# RNA-guided RNA silencing by an Asgard archaeal Argonaute

**DOI:** 10.1101/2023.12.14.571608

**Authors:** Carolien Bastiaanssen, Pilar Bobadilla Ugarte, Kijun Kim, Yanlei Feng, Giada Finocchio, Todd A. Anzelon, Stephan Köstlbacher, Daniel Tamarit, Thijs J. G. Ettema, Martin Jinek, Ian J. MacRae, Chirlmin Joo, Daan C. Swarts, Fabai Wu

## Abstract

Eukaryotic Argonaute proteins achieve gene repression and defense against viruses and transposons by RNA-guided RNA silencing. By contrast, known prokaryotic Argonautes adopt single-stranded DNA as guides and/or targets, leaving the evolutionary origin of RNA-guided RNA silencing elusive. Here, we show an evolutionary expansion of Asgard archaeal Argonautes (asAgos), including the discovery of HrAgo1 from the Lokiarchaeon ‘*Candidatus* Harpocratesius repetitus’ that shares a common origin with eukaryotic PIWI proteins. HrAgo1 exhibits RNA-guided RNA cleavage *in vitro* and RNA silencing in human cells. The cryo-EM structure of HrAgo1 combined with quantitative single-molecule experiments reveals that HrAgo1 possesses hybrid structural features and target binding modes bridging those of the eukaryotic AGO and PIWI clades. Finally, genomic evidence suggests that eukaryotic Dicer-like processing of double-stranded RNA likely emerged as a mechanism of generating guide RNA for asAgos prior to eukaryogenesis. Our study provides new insights into the evolutionary origin and plasticity of Argonaute-based RNA silencing.

## Main

Argonaute proteins facilitate guide oligonucleotide-mediated binding of nucleic acid targets to perform a wide range of functions in prokaryotes and eukaryotes. In eukaryotic RNA silencing pathways, sequence-specific repression of target RNAs is achieved by Argonaute proteins loaded with small guide RNAs^1–5^. Canonical eukaryotic Argonautes (eAgos) can be subdivided into two clades, AGO and PIWI, which are distributed broadly, albeit heterogeneously, across eukaryotic lineages^6^. AGOs and PIWIs are strictly conserved and arguably best studied in Metazoa (animals), where they rely on various guide generation pathways and carry out distinct physiological functions. Metazoan AGOs use small interfering RNA (siRNA) and/or microRNA (miRNA) guides, generated from double-stranded RNA (dsRNA) by Dicer-like RNase III family proteins, to post-transcriptionally regulate gene expression^7,8^. In general, base-pairing of a short region at the 5’ end of miRNAs termed the ‘seed’ (nucleotides 2-8) to a target RNA is sufficient for AGOs to bind target RNA^8^. By contrast, PIWIs generally show lower seed binding strength and target RNA binding requires extended base-pairing in the central region of the guide to achieve stable binding^5^. Additionally, metazoan PIWI-interacting RNA (piRNA) guides are generated from longer single-stranded RNA by Zucchini, to suppress transposable elements (TEs)^9^. Both the arms race against TEs and global gene silencing are critical drivers of eukaryotic genome evolution^10–15^. As such, the origin and differentiation of AGO and PIWI have broad implications for the emergence and expansion of the eukaryotic tree of life. However, it is unclear how the divergence between AGO- and PIWI-based RNA silencing pathways originated and whether they have consistent signatures across the expansive eukaryotic tree of life.

Prokaryotic Argonautes (pAgos) are a highly diverse protein family with functions ranging from prokaryotic immunity by neutralizing foreign DNA^16–19^ or inducing cell death in invaded cells (abortive infection)^20,21^, to aiding in genome replication and recombination^22,23^. All pAgos characterized to date interact with DNA guides and/or targets; no known pAgo exclusively facilitates eAgo-like guide RNA-mediated RNA targeting. Thermophilic euryarchaeal Argonautes, which have previously been suggested to be most closely related to eukaryotic Argonautes^24^, exclusively mediate DNA-guided targeting of invading DNA^19,25^. Furthermore, no dedicated guide RNA-generating systems, such as homologs of eukaryotic Dicer or Zucchini, have been found associated with pAgos. Hence, with these apparent mechanistic differences between pAgos and eAgos, it was thought that RNA-guided RNA-targeting Argonautes, along with their associated guide RNA-generating pathways, have arisen after eukaryogenesis and before the last eukaryotic common ancestor (LECA)^10,26^. Here we show that an Asgard archaeal Argonaute mediates RNA-guided RNA silencing, providing new insights into the origin and diversification of eukaryotic RNA silencing pathways.

### Asgard archaeal diversification gave rise to eAgo-like Argonautes

Eukaryotes are thought to have evolved from an archaeon belonging to Asgard archaea (or Asgardarchaeota)^13,27–30^. We thus set out to explore the presence of Argonaute proteins in these organisms using a custom hidden Markov model based on the conserved MID-PIWI domains (See Methods). In 496 available Asgard archaeal metagenome-assembled genomes (MAGs), we identified a total of 138 Asgard archaeal Argonaute sequences (asAgos, see Table S1). Maximum-likelihood phylogenetic analysis shows that asAgos are polyphyletically distributed over 15 subclades located across the phylogenetic tree of Argonaute proteins, including subclades 1, 11, and 13 that respectively appear basal to the previously classified long-A pAgos, long-B pAgos, and short pAgos^20,21,31^ (Fig. 1a). Like many other prokaryotic defense systems^32^, pAgos are present only in a fraction of prokaryotes. Here we found Argonaute-encoding genes in 21.5% (83/387) of the quality-filtered Asgard archaeal MAGs, higher than any other prokaryotic phylum as classified by the Genome Taxonomic Database^33^ (GTDB v207). This apparent gene enrichment and diversification imply that Asgard archaea may have adopted Argonaute proteins for diversified functions (Fig. 1b).

**Figure 1.**
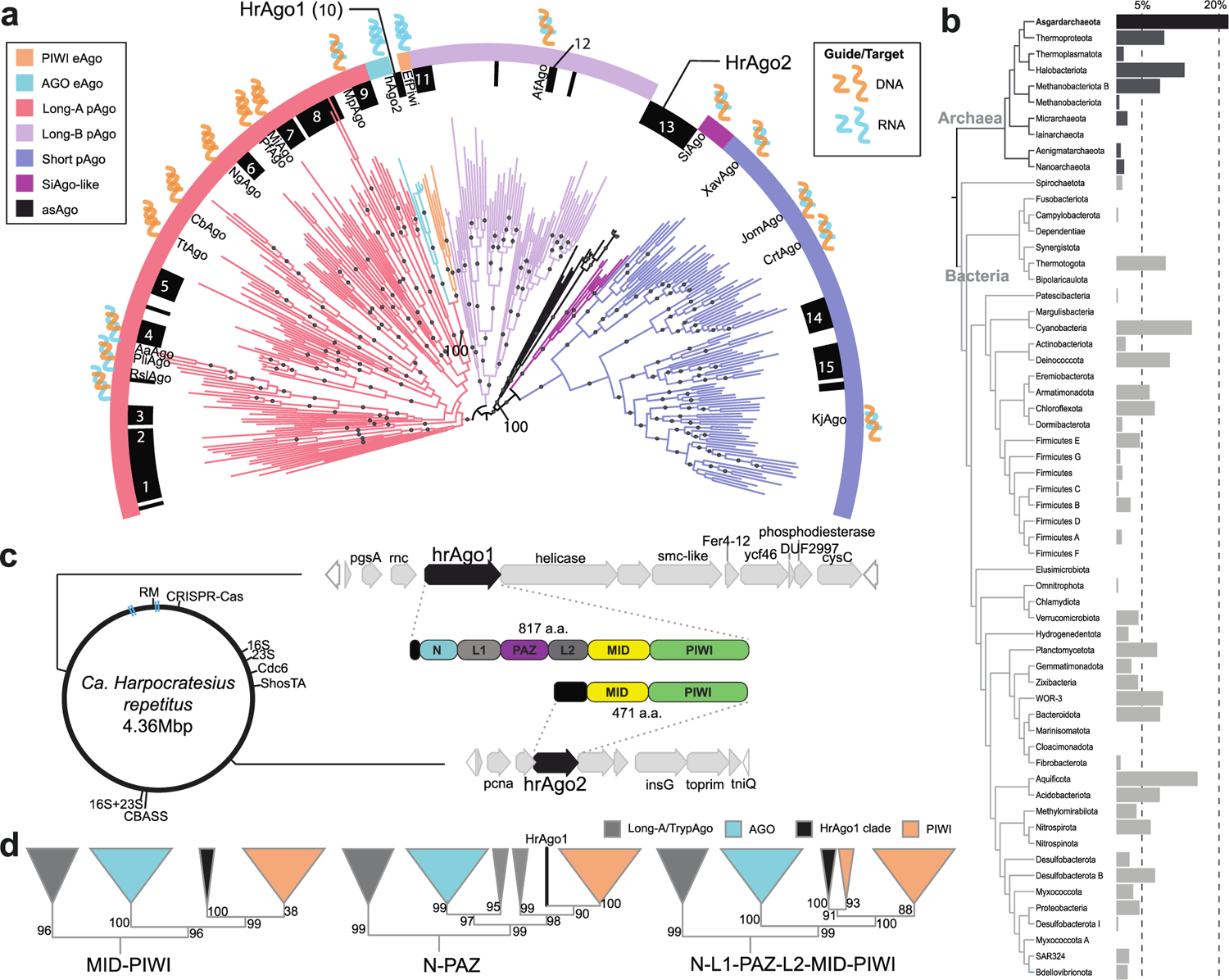
The expanded Argonaute diversity in Asgard archaea. **a**, Maximum-likelihood phylogenetic analysis of the MID-PIWI domains of Argonaute proteins showing that asAgos are polyphyletic (black pallets, subclades 1-15 denoted). 334 representative sequences and 572 sites were analyzed using IQ-tree based on Q.pfam+C60+F+G4 model. Different branch and ring colors indicate different major Argonaute clades. Various representative Argonautes (see Methods) and their primary guide/target preferences are indicated, while they may have secondary guide/target use. Ultrafast bootstrap 2 (UFBoot2) values above 95, calculated based on 1000 replicates, are shown in black circles. HrAgo1 and HrAgo2, and the UFBoot2 values at the base of their respective clades are highlighted. **b**, Fraction of Argonaute-encoding genomes in different prokaryotic phyla. **c**, Genomic depiction of Asgard archaeon ‘*Ca.* H. repetitus’, where the genes encoding 16S and 23S rRNA, origin of replication protein Cdc6, and putative immune systems are indicated. The synteny and predicted domain compositions of genes surrounding pAgo-encoding genes are highlighted. RM, restriction-modification system. Blue bars indicate two genome assembly gaps with undetermined sequences. **d**, Maximum-likelihood phylogenetic analysis of AGO, PIWI, and HrAgo1 using different domain combinations (indicated at the bottom) illustrates the robust position of HrAgo1 basal to the PIWI clade. UFBoot2 values calculated based on 1000 replicates are indicated.

Strikingly, we found that ‘*Candidatus* Harpocratesius repetitus FW102’^13^, a deep-sea rock-dwelling Lokiarchaeia archaeon named after the Greek god of silence, encodes two asAgos belonging to Asgard-specific-clades distinct from known pAgos. The HrAgo1 subclade clusters with eAgos while the HrAgo2 subclade comprises a mixture of long and short asAgos basal to all short pAgos (Fig. 1a). Both are encoded in operon-like gene clusters outside of other genomic defense islands, including CRISPR-Cas and CBASS systems. The HrAgo2 operon encodes various components involved in transposition (InsG and TniQ) and DNA replication (PCNA and TOPRIM). This is different from known short pAgos or SiAgo-like pseudo-short pAgos, which cooperate with immune effectors encoded in their gene neighborhoods to trigger cell death^20,21^. The HrAgo1 operon is also unique in that flanking *hrAgo1* are an *rnc* gene, encoding a protein that comprises an RNaseIII domain and a double stranded RNA binding domain (dsRBD), and a gene encoding a HEDxD/H helicase (Fig. 1c). Both proteins share functional domains with eukaryotic Dicer enzymes involved in guide RNA biogenesis.

HrAgo1 shows higher similarity to well-studied PIWIs (25-27% sequence identity) and AGOs (23-24%), than to various other pAgos (16-21%) (Fig. S1). To date, the only eAgo-like HrAgo1 homolog that we could identify is a truncated asAgo sequence found in a Lokiarchaeon assembled from a Siberian soda lake metagenome^34^, with 34% sequence identity to HrAgo1 across the obtained L2-MID-PIWI segment. Although a subclade of asAgos (subclade 9 in Fig. 1a) appeared to be basal to the whole eAgos clade in this analysis, the inferred evolutionary relation is supported by a low bootstrap value and unstable against changes in phylogenetic methods (Fig. S2). To further elucidate the relation between HrAgo1 and eAgos, we expanded the sampling of AGO and PIWI clade homologs across the eukaryotic tree of life and performed Maximum Likelihood analyses using above-found Long-A pAgos/asAgos and the non-canonical Trypanosome-specific TrypAgos as outgroup. When analyzed using the conserved MID-PIWI domains commonly used for Argonaute phylogeny^24^, we found that the HrAgo1 clade is positioned as sister group to the PIWI clade (Fig. 1d). Additionally, we examined the more variable N-L1-PAZ domains as well as the full-length N-L1-PAZ-L2-MID-PIWI domains, which, despite a few unstable branches of pAgos and eAgos whose evolutionary positions are uncertain, further confirmed the monophyly of HrAgo1 as being basal to the PIWI clade (Fig. 1d, Fig. S3). Combined with phylogenomic studies supporting an asgard archaeal origin of eukaryotes, our data suggest that eukaryotic PIWIs and HrAgo1 evolved from a common ancestor, prompting us to study the molecular mechanism and function of HrAgo1.

### HrAgo1 mediates RNA-guided RNA cleavage

The most apparent differences between eAgos and pAgos are their guide and target preferences. We thus analyzed the oligonucleotides that associate with HrAgo1 upon heterologous expression in *E. coli*. 5’-end ^32^P-labelling of the associated nucleic acids reveals that HrAgo1 associated with 15-25 nt-long small RNAs, but not with DNA (Fig. 2a). Corroborating the ^32^P-labeling-based detection, small RNA sequencing analysis confirmed that HrAgo1-associated small RNAs are mostly 15-25 nt in length (Fig. 2b). The small RNAs have a bias for uracil (U) at their 5’ end (65%), similar to the guide 5’-end preference observed for most examined PIWIs and AGOs^9,35^. Furthermore, a bias for U is observed to a lesser extent at position 2 (47%) and 3 (49%) of the guide RNA (Fig. 2c). Since previous studies have shown that nucleic acids co-purified with heterologously expressed pAgos generally match the types of their naturally preferred guides^16–18,20,36^, our data thus suggest that HrAgo1 utilizes guide RNAs, akin to eAgos.

**Figure 2.**
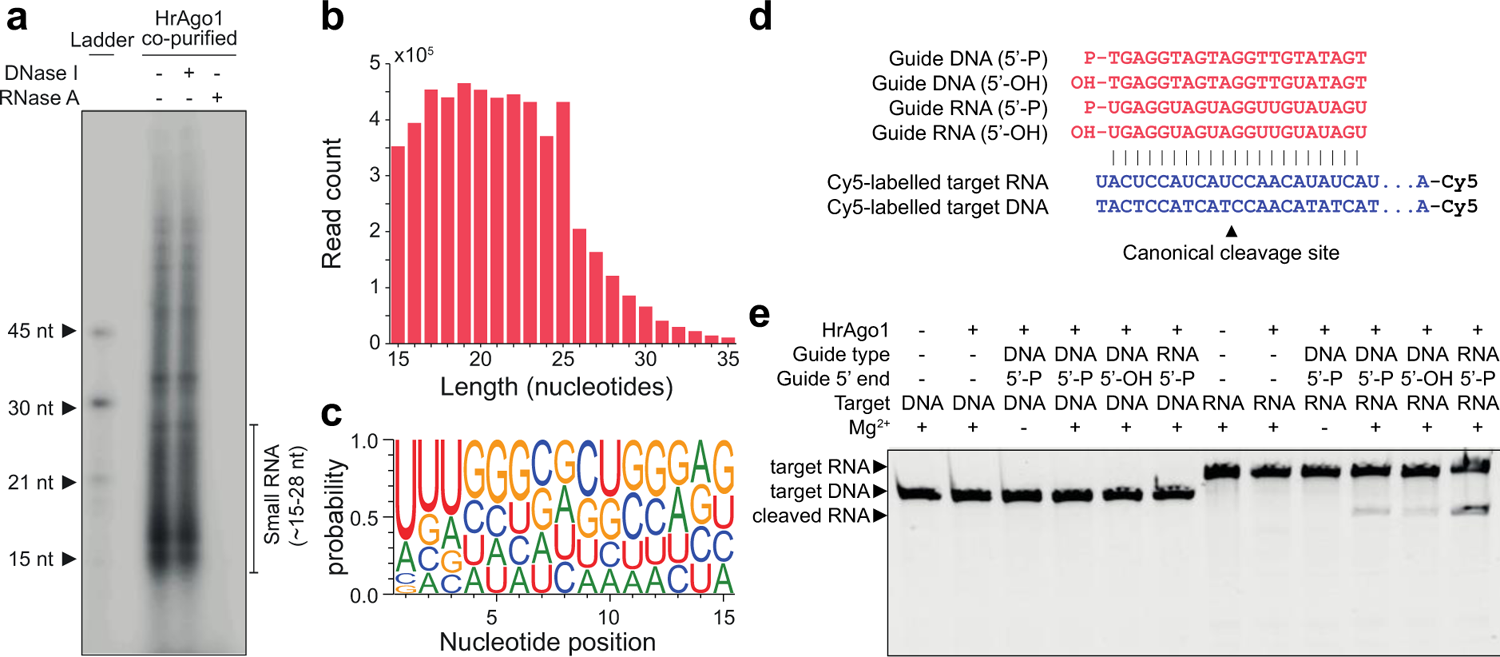
HrAgo1 mediates RNA-guided RNA cleavage. **a**, HrAgo1 associates with 5’ phosphorylated (5’ P) small RNAs *in vivo*. Nucleic acids that co-purified with HrAgo1 were [γ-^32^P] labeled, treated with RNase A or DNase I, and resolved on a denaturing gel (15% polyacrylamide 7M urea). nt: nucleotides. **b**, Length distribution of small RNAs associated with HrAgo1 as determined by small RNA sequencing. **c**, Small RNAs associated with HrAgo1 have a bias for uracil bases at the 5’ end. **d**, Sequences of guide and target oligonucleotides used in *in vitro* cleavage assays. **e**, HrAgo1 cleaves ssRNA (but not ssDNA) targets with ssRNA guides, and ssDNA guides at lower efficiency, in the presence of Mg^2+^. HrAgo1 was incubated with ssDNA or ssRNA guides and Cy5-labelled ssDNA or RNA targets. Cy5-labeled cleavage products were resolved through denaturing (7M urea) polyacrylamide gel electrophoresis and visualized by fluorescence imaging.

Next, we analyzed HrAgo1 guide/target preferences *in vitro*. Upon incubation of HrAgo1 with 21-nt single-stranded (ss)DNA or ssRNA guide oligonucleotides and complementary 5’ Cy5-labelled ssDNA or ssRNA targets (Fig. 2d, Table S2), HrAgo1 demonstrated ssRNA-guided cleavage of RNA targets in a magnesium-dependent manner, while it was unable to cleave DNA targets (Fig. 2e). Of note, guide ssDNAs also facilitated cleavage of RNA targets, but with lower efficiency compared to guide ssRNAs (Fig. 2e), similar to the *in vitro* behavior of human AGO2 (hAgo2)^37^. Combined, these results show that, compared to other known pAgos, the prokaryotic HrAgo1 mechanistically acts more similarly to RNA-guided RNA-targeting eAgos.

### Structural architecture of HrAgo1

To illuminate the structural basis for RNA-guided RNA cleavage by HrAgo1, we examined HrAgo1 in complex with a 21-nucleotide guide RNA by cryogenic electron microscopy (cryo-EM) and single particle analysis. The resulting reconstruction, determined at a resolution of 3.4 Å, reveals a binary HrAgo1-guide RNA complex (Fig. 3a-d, Fig. S4, Table S3). Resembling eAgos and long pAgos, HrAgo1 adopts a bilobed conformation in which one lobe comprises the N-terminal, linker L1, PAZ, and linker L2 domains, connected to the second lobe comprised of the MID and PIWI domains (Fig. 3c-d). The first six nucleotides of the guide RNA 5’ end (g1-g6) are ordered in the cryo-EM map (Fig. 3b-d). Low resolution density for four nucleotides at the 3’ end of the guide RNA (g18-g21) is also apparent but uninterpretable, while the remainder of the guide RNA is unstructured (Fig. 3b-d). In accordance with its phylogeny, an all-against-all comparison^38^ of experimentally determined structures of Argonaute-family proteins positions HrAgo1 between pAgos and eAgos, and closest to the PIWI-clade Siwi (Fig. 3e).

**Fig. 3:**
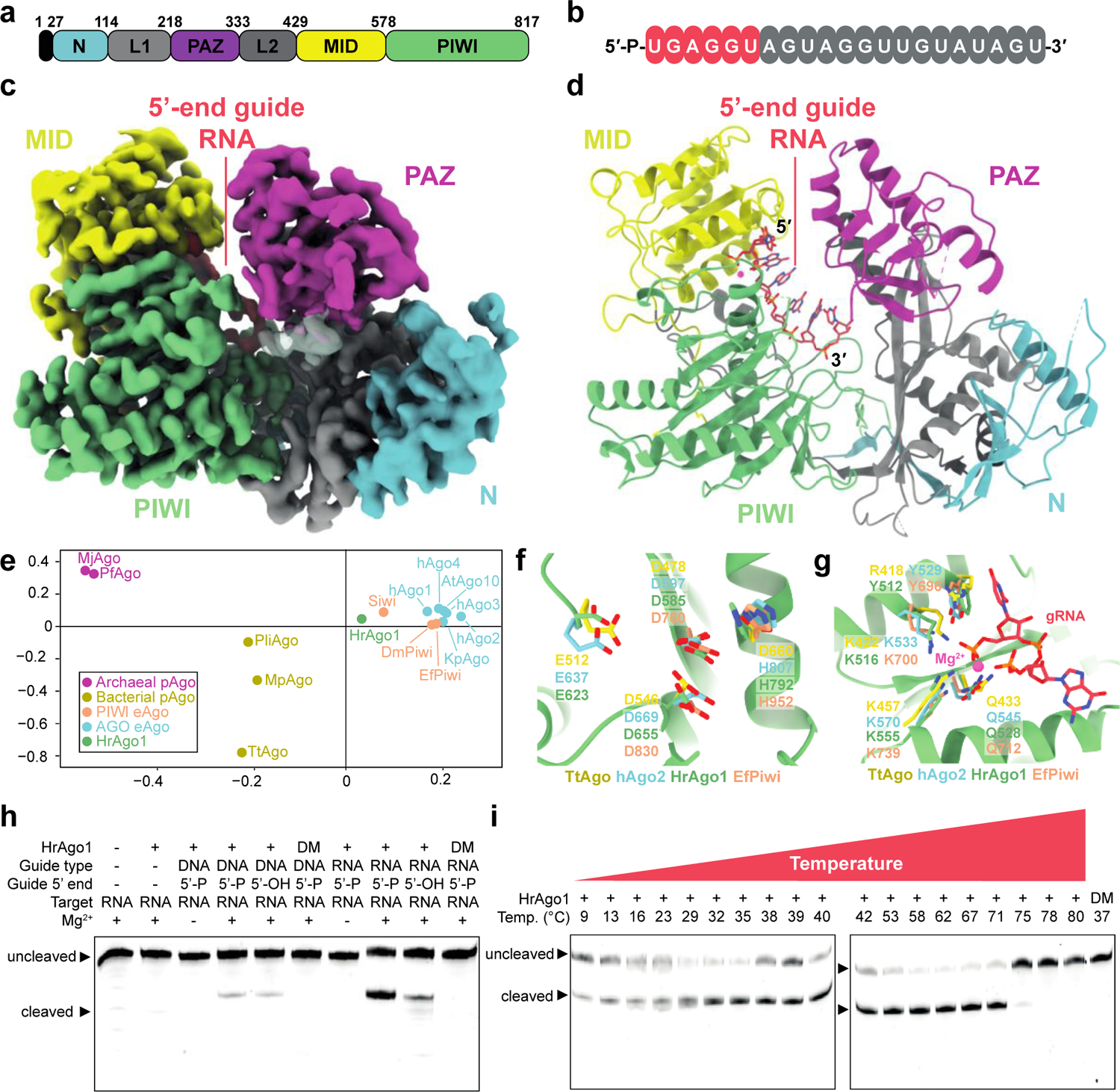
Molecular architecture of HrAgo1 bound to a guide RNA. **a**. Schematic diagram of the domain organization of HrAgo1. N, N-terminal domain; L1 and L2, linker domains; PAZ, PIWI-ARGONAUTE-ZWILLE domain; MID, Middle domain; PIWI, P-element induces wimpy testis domain. **b**, Schematic representation the HrAgo1-bound guide RNA. Structurally ordered residues are colored red, while disordered residues are colored grey. **c**, Cryo-electron microscopic (cryo-EM) density map of HrAgo1 bound to a guide RNA Cartoon, colored according to individual domains. The unmodeled 3’-end guide RNA density is represented as a transparent surface. **d**, Cartoon representation of the overall structure of the HrAgo1-guide RNA complex. **e**, All-against-all structure comparison of selected Argonaute proteins. **f**, Close-up view of the HrAgo1 catalytic site aligned to that of other representative Argonaute proteins. **g**, Close-up view of the HrAgo1 guide RNA 5’-end binding site in the MID domain aligned to that of other representative Argonaute proteins. **h**, Efficient HrAgo1-mediated RNA cleavage requires a guide RNA with a 5’ phosphate and an intact catalytic site. HrAgo1 was incubated with ssDNA or ssRNA guides and Cy5-labelled ssRNA targets. DM: HrAgo1 catalytic mutant with D585A and E623A substitutions. **i**, HrAgo1 mediates RNA-guided RNA cleavage at temperatures ranging from 9 °C to 71 °C. HrAgo1 was incubated with ssRNA guides and Cy5-labelled ssRNA targets. For h and i, Cy5-labelled cleavage products were resolved on a denaturing (7M urea) polyacrylamide gel and visualized by fluorescence imaging.

The catalytic tetrad of HrAgo1 comprises residues Asp585, Glu623, Asp655, and His792 (Fig. 3f). In the structure, all four catalytic residues are ordered and in position to mediate divalent cation binding and catalysis, akin to the catalytic site of AGO structures^2^. This implies that HrAgo1 adopts a catalytically active conformation. The 5′-terminal phosphate group of the guide RNA is sequestered in the MID domain binding pocket through interactions with residues (Phe512, Lys516, Asn528, and Lys555) that are conserved in most Argonautes^39^ (Fig. 3g). The negative charge of two phosphates of guide RNA nucleotides 1 and 3, as well as that of the C-terminal carboxyl group of HrAgo1, are neutralized by a Mg^2+^ ion as is observed in pAgos and PIWIs (Fig. 3g). Instead of Mg^2+^, Metazoan AGOs use another lysine residue in this pocket^40^. A catalytic double mutant (D585A & E623A, HrAgo1^DM^) did not mediate RNA cleavage, confirming that the catalytic DEDH motif in the PIWI domain facilitates target cleavage (Fig. 3h). Corroborating the observed interactions with the 5’-phosphate, HrAgo1 showed higher activity with guide RNAs that are 5’-phosphorylated compared to guide RNAs with a 5’-hydroxyl group (Fig. 3h). Remarkably, HrAgo1 mediated RNA-guided RNA cleavage at temperatures ranging from 9 °C to 71 °C (Fig. 3i), coinciding with a steep temperature gradient around the hot hydrothermal vents that ‘*Ca.* H. repetitus’ resided. Such an extraordinarily broad temperature adaptation apparently places HrAgo1 between the temperature ranges of mesophilic eAgos with those from the euryarchaeal pAgos, which mostly function at temperatures above 75 °C^19,41,42^. Our structural data combined with biochemical experiments thus illuminate the mechanistic adaptation of the archaeal HrAgo1 as an eAgo-like RNA-guided RNA-cleaving enzyme.

### HrAgo1 structure bridges AGO and PIWI

The ability to rapidly find complementary mRNA targets in a crowded cellular environment and to effectively distinguish them from other transcripts is critical for the functions of eAgos. Argonaute proteins achieve efficient target binding by ordering the ‘seed’ region of the guide strand (nucleotides 2-8) in an A-form helical conformation^2,43,44^. AGO-clade eAgos achieve full seed pre-organization using a loop that binds the g5-g6 backbone and a helix-7 which organizes g7–g8^45^, facilitating strong target association at short matching lengths^46^. By contrast, known PIWI-clade eAgos only pre-organize the first few nucleotides of the seed (typically g2-g4), and require further guide-target pairing beyond the seed to achieve stable target binding^3,5^. In HrAgo1, guide RNA nucleotides g2-g6 are pre-ordered in an A-form-like helical conformation (Fig 4a). Remarkably, HrAgo1 simultaneously possesses structural features related to guide RNA ordering that are typically observed in either AGOs or in PIWIs (Fig. 4b). HrAgo1 is AGO-like in that it uses contacts in the g5-g6-binding loop (H738 and the main chain amide of R746) to organize guide nucleotides g5–g6, and is also PIWI-like in that helix-7 is tilted away from the seed, leaving g7–g8 disordered. Tilting of helix-7 in HrAgo1 may be attributed to the presence of a loose “seed-gate” structure (residues Phe333-Gln354), also present typically found in PIWIs, but more stretched in AGOs^3,5^. In PIWIs, the seed-gate structure has been proposed to enable extended target probing at positions g5-g9^5^. Overall, HrAgo1 structure pre-organizes guide RNA in a manner more similar to AGOs, while showing features similar to PIWIs that may influence its target binding.

**Fig. 4:**
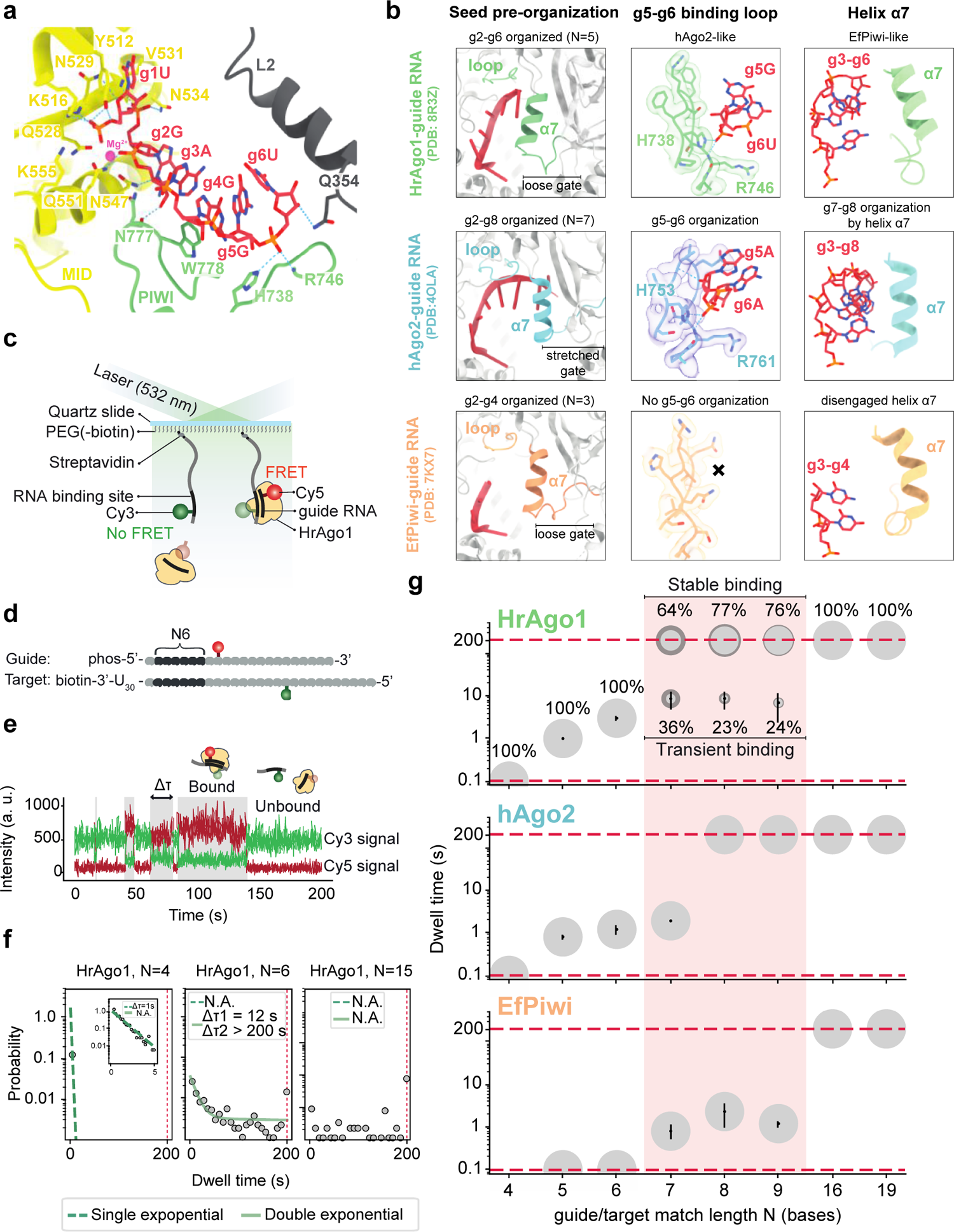
HrAgo1 displays a unique hybrid mode of guide organization and target binding. **a**, Close-up view of guide RNA organization by HrAgo1. **b**, Comparison of structural features involved in guide RNA seed segment organization in HrAgo1, EfPiwi, and hAgo2. **c**, Schematic of the single-molecule binding assay. Only when the HrAgo1-guide complex binds to the target, FRET will occur. **d,** Schematic representation of a guide and target used in the single-molecule binding assay. Complementary nucleotides are indicated in dark grey and mismatched nucleotides are shown in light grey. N6 indicates base pairing with nucleotides 2-7 of the guide. **e**, A representative time trace with four binding events, of which the dwell time (Δτ) of one is indicated. **f,** Dwell time distributions for N4, N6 and N15. N4 and N6 are best fit with a single and double exponential, respectively. N15 cannot be fit and shows stable binding. The distributions and fits for the other match lengths and representative time traces can be found in Fig. S5. **g,** Bubble plots showing the increase of dwell times for increasing complementarity between the guide and target for HrAgo1, EfPiwi and hAgo2. The area of the bubbles corresponds to the percentage of the total population belonging to this sub-population. The dashed lines indicate the time resolution (0.1 s) and the observation time limit (200 s). Error bars and the darker shaded area of the bubbles indicate the standard deviation of at least three independent experiments. The dwell times for EfPiwi were obtained in a similar way as for HrAgo1. For hAgo2, previously published dwell times were used^46^.

### HrAgo1 displays a unique hybrid mode of target binding

To investigate the target RNA binding kinetics of HrAgo1, we performed a single-molecule fluorescence resonance energy transfer (FRET) binding assay (Fig. 4c). Guide and target RNAs were labeled with Cy5 and Cy3 dyes respectively so that binding of the HrAgo1-guide complex to the target gives rise to a high FRET signal (Fig. 4d-f, Fig. S5a, Table S4). We quantitatively investigated the binding of the HrAgo1-guide RNA complex to target RNAs with varying guide-target complementarity and compared that to the same experiments performed with EfPiwi and to hAgo2 data from literature^46^ (Fig. 4g, Fig. S6). The interactions between HrAgo1 and the target became observable when the latter matches the nt 2-4 (N3) positions of the guide RNA, and the dwell time increases drastically with the increase in guide-target match length (Fig. 4f and 4g, Fig. S5). At these short match lengths, the dwell time distribution follows a simple exponential decay, similar to previous observations of hAgo2^46^. Starting from N6, the majority of the guide-target association events of HrAgo1 and hAgo2 persist beyond the experimental time limit of 200 s (Fig. 4g). The overall binding kinetics of HrAgo1 are thus similar to the behavior of hAgo2. This contrasts EfPiwi, which only shows observable interactions with the target at a match length of N6, and shows stable binding only at N15, in agreement with structural predictions^5^.

While HrAgo1 facilitates prolonged binding for most of the guide-target pairs between N6 and N8, a notable sub-population remains only transiently bound, resembling the behavior of EfPiwi (Fig. 4g). The appearance of a second population has been occasionally observed previously when the binding pocket of Argonaute interacts with a specific species of nucleotide in the first position of the target, e.g. deoxyguanosine by TtAgo^47^ and deoxyadenosine by hAgo2^48^. However, the two-population behavior we observe here is independent of the identity of the first target nucleotide (Fig. S5b-c), suggesting that HrAgo1 intrinsically utilizes two modes of target search, i.e. an overall strong seed binding mode as observed for AGOs, and a second mode of transient seed binding akin to PIWIs. Consistent with its hybrid structural features, HrAgo1 thus facilitates a unique hybrid mode of guide RNA-mediated target RNA binding.

### HrAgo1 mediates RNA silencing in human cells

The physiological function of HrAgo1 can provide clues to the emergence and diversification of RNA silencing pathways. However, Asgard archaea are notoriously slow-growing, largely uncultivated, and not genetically accessible. Furthermore, ‘*Ca.* H. repetitus’ was enriched from undetectable to only 1% of the community on a low-biomass hydrothermal rock^13^, and is therefore not a suitable host for physiological characterization of HrAgo1. Given the structural and mechanistic resemblance of HrAgo1 to eAgos, particularly its main binding characteristics resembling that of the human hAgo2, we examined whether HrAgo1 can perform RNA silencing in a human cell line.

To exclude any endogenous RNA interference (RNAi) activity, we adopted an HCT116 cell line in which *hAgo1/2/3* genes are knocked out (*AGO1/2/3* KO HCT116)^49^. We first performed stable transfection of pLKO.1 puro-pri-mir-1-1 vector, which encodes puromycin *N*-acetyltransferase that confers resistance to puromycin and a primary hairpin transcript (pri-mir-1-1) that acts as a precursor for mature miR-1-1 whose expression is suppressed in the parental cells^50^ (Fig. 5a, Fig. S7a). Puromycin-selected cells were then co-transfected with a dual-expression vector that encodes firefly luciferase (Fluc) and *Renilla* luciferase (Rluc), as well as with an expression vector encoding FLAG-tagged HrAgo1 (FLAG-HrAgo1) (Fig. 5b, Fig. S7b). In addition, vectors expressing superfolder GFP (sfGFP) and FLAG-tagged hAgo2 (FLAG-hAgo2) were used as negative and positive controls, respectively. The 3′ UTR of the Fluc gene has two binding sites with perfect complementarity to miR-1-1, which allows Ago-mediated silencing of Fluc expression. To monitor miR-1-1-guided Fluc silencing, we performed qPCR to measure the relative expression level between target Fluc mRNA and the control Rluc mRNA. Remarkably, cells in which miR-1-1 and HrAgo1 were co-expressed, showed a significant (p<0.01) decrease in the Fluc/Rluc mRNA ratio compared to cells in which the sfGFP control was co-expressed with miR-1-1 (Fig. 5c). Moreover, the level of post-transcriptional repression by HrAgo1 was comparable to that of the hAgo2 without significant difference. This demonstrates that HrAgo1 is capable of RNA silencing in human cells. Furthermore, the use of a dsRNA hairpin precursor to supply guide RNAs suggests that HrAgo1 can accommodate guide RNAs generated by the canonical miRNA biogenesis pathway involving the Microprocessor complex and Dicer.

**Figure 5.**
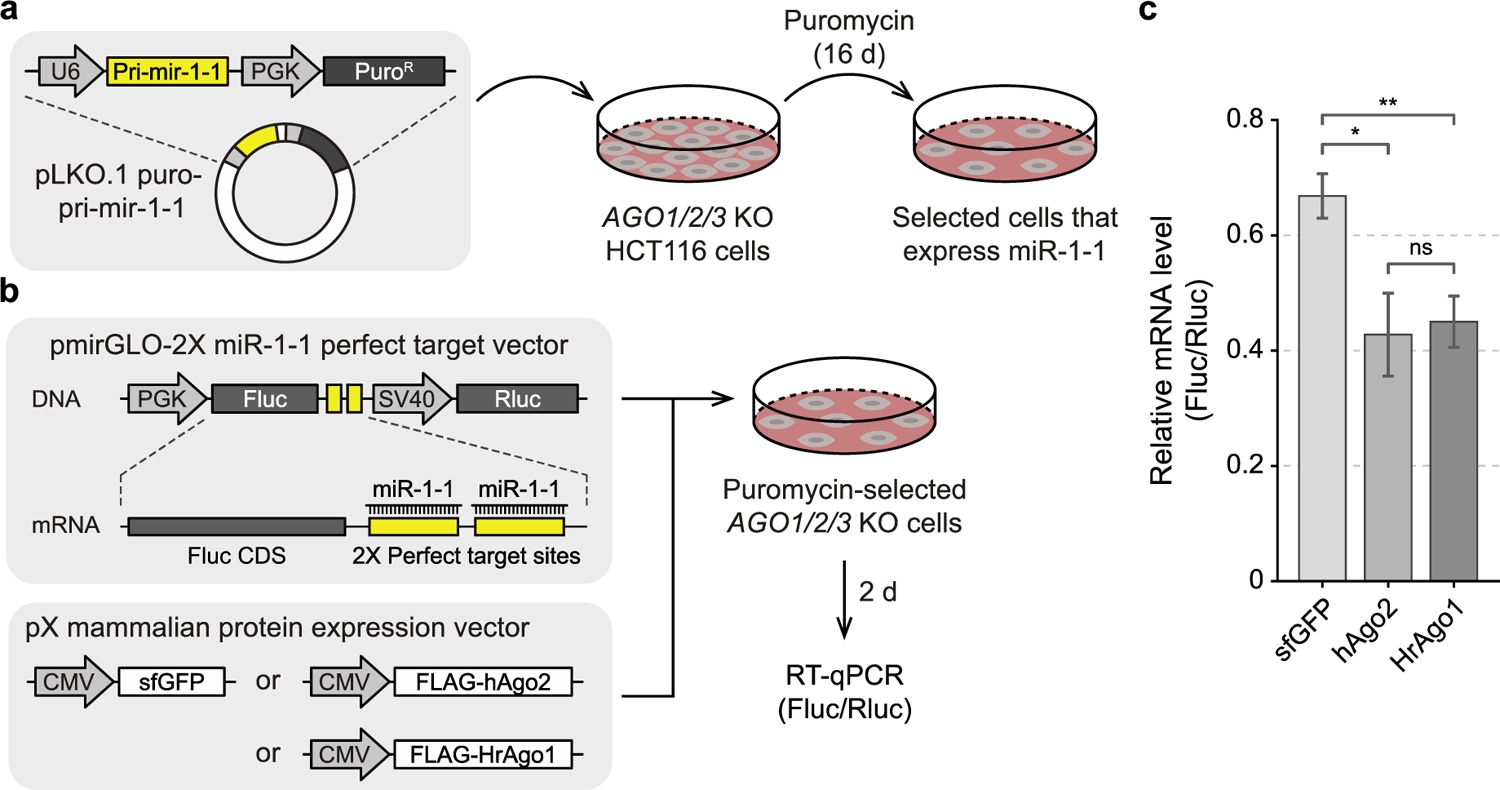
HrAgo1 mediates RNA silencing in human cells. **a**, A schematic diagram for stable transfection of miR-1-1. pLKO.1 puro-pri-mir-1-1 vector was transfected into *AGO1/2/3 KO* HCT116 cells, which were subsequently subjected to puromycin selection for 16 days to generate cells that stably express miR-1-1. **b,** A schematic diagram for the RNAi rescue experiment. Puromycin-selected cells were co-transfected with a dual-luciferase expression vector containing two perfect target sites for miR-1-1 in the 3′ UTR of the firefly luciferase gene (Fluc) and a protein expression vector encoding sfGFP or hAgo2 or HrAgo1. 2 days after the transfection, total RNA was isolated and subjected to RT-qPCR. **c,** qPCR results for relative mRNA expression levels between firefly luciferase (Fluc) and *Renilla* luciferase (Rluc). Bars indicate mean ± SD (n=3, biological replicates). ns, not significant; **, p<0.01; *, p<0.05 by independent samples t-test.

## Discussion

In this study, we explored the diversity of Asgard archaeal Argonautes and characterized HrAgo1, an eAgo-related asAgo. This extends our understanding on the evolutionary origin and diversification of pAgos as well as their relation with eAgos. Resolving long-term evolutionary trajectory is challenging, especially considering that a defense-related gene like *pAgos* could have potentially undergone gain, loss, and horizontal gene transfer (HGT) at a high frequency^51^. Our global phylogenetic analysis shows that asAgos exhibit a striking diversity including new, deep-branching subclades basal to Long-A, Long-B, and short pAgos. Asgard archaea may thus have been the donors of pAgos HGT for various bacterial and archaeal lineages, which adopted different types of guide and target nucleic acids^16–23,25^. The molecular basis underlying the differences in guide/target specificity among pAgos, as well as between pAgos and eAgos, is yet unclear. Studying the structural and biochemical properties of these deep-branching asAgos can provide valuable insights into the origin and diversification of pAgos.

We found that HrAgo1 is a prokaryotic Argonaute capable of RNA-guided RNA silencing, consistent with its phylogenetic position basal to the PIWI-clade eAgos. These properties of HrAgo1 also provided a unique opportunity for us to gain insights into the diversification of AGO and PIWI at the molecular level. Based on our comparative analyses of structural and single-molecule FRET data between HrAgo1 and different eAgos, we hypothesize that the common ancestor of AGO- and PIWI-clade eAgos had a g5-g6 pre-organizing loop, while its helix-7 did not embrace g7-g8. AGOs kept and further refined the g5-g6 organizing loop, while repositioning helix-7 to enable g7-g8 pre-organization, allowing strong target association at short matching lengths to facilitate post-transcriptional silencing of a multitude of genes^2,7,8,21,46^. The structure of HrAgo1 suggests that early-branching PIWIs kept the g5-g6 loop, while later-evolved PIWIs lost it, giving rise to the more relaxed targeting preferences of metazoan piRNAs that enable defense against evolving genomic threats^5,52^.

As HrAgo1 is positioned basal to the PIWI subclade while no stable sister clade of the broader eAgo clade has been identified, the exact evolutionary origin of eAgos and their associated pathways remains to be resolved. Phylogenetic analyses suggested that some candidate asAgos (e.g. subclade 9 in Fig. 1a, also see Fig. S2) may be the closest relatives to all eAgos, though with weak phylogenetic support. Another defining feature of eAgos that differ from all characterized pAgos so far is their associations with dedicated RNA guide generation mechanisms, such as Dicer-based dsRNA processing pathway known to provide guides for AGOs^53^ and Zucchini-based ssRNA processing pathway for metazoan PIWIs^9^. We found that *hrAgo1* is flanked by genes encoding RNaseIII/dsRBD domain-containing Rnc and DExD/H domain-containing Helicase (Fig. 1b). These domains are found in the eukaryotic Dicer and Dicer-like proteins^54–57^, which suggests that the neighboring genes of hrAgo1 may adopt a guide-generating system that processes dsRNA. While the molecular mechanisms of these proteins require further experimental validation, we did find multiple examples of *rnc-asAgo-helicase* gene associations (Fig. 6a); additional associations may have been missed due to fragmented genome assembly. Notably, syntenic associations where Rnc and asAgos are consecutively encoded on the same DNA strand are only found in the HrAgo1 clade and asAgo subclade 9 phylogenetically close to eAgos, which provides an additional support for their close relation to eAgos (Fig. 6a).

**Figure 6:**
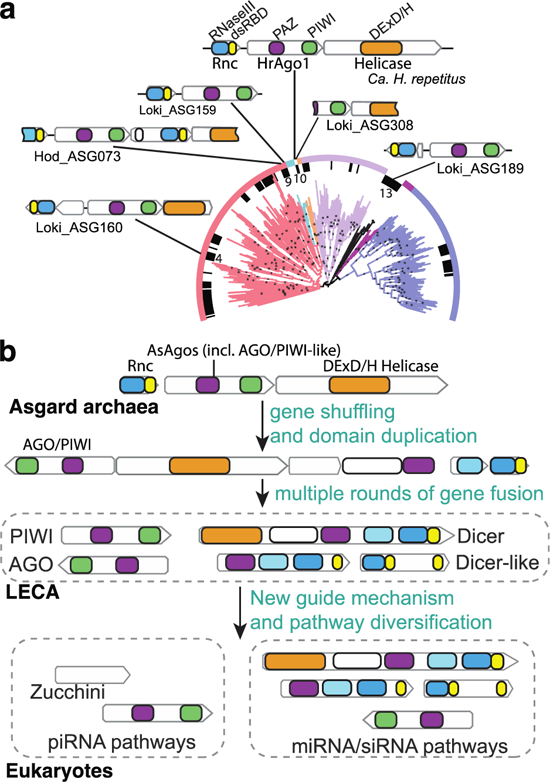
Origin and diversification of the eukaryotic RNA silencing pathways. **a**, Multiple *rnc-asAgo-helicase* gene clusters distributed across the phylogenetic tree of Argonaute. The different domains are indicated by different colors as depicted on the top. Lines link the gene schematics with the phylogenetic positions of the corresponding asAgo. Numbers on the inner circle denote asAgo subclades as defined in Fig. 1a. **b**, A hypothetical model of the emergence of canonical Dicer as well as the RNA silencing pathways through gene fusion from ancestral gene clusters containing genes encoding Rnc, asAgo, and DExD/H Helicase. Other related components, such RNA-dependent RNA polymerase, of the pathways are omitted in the illustration.

Combined, our analyses provided new clues to the origin of eukaryotic RNA silencing pathways. Previous pan-eukaryote analyses have implicated that LECA likely encoded RNA silencing machineries comprising an AGO, a PIWI, a Dicer, as well as an RNA-directed RNA polymerase (RdRp), which partners with Dicer in some physiological contexts^6^. The evolutionary paths leading to the emergence of such an RNA silencing pathway in LECA was, however, unclear. Existing models posited that RNA silencing most likely emerged after eukaryogenesis based on two main observations: 1) archaeal pAgos previously found to be closest to eAgos performed DNA-guided DNA cleavage, contrasting the RNA-guided RNA cleaving eAgos, 2) the RNaseIII domains and Helicase domains of Dicer appeared to have respectively originated in bacteria and archaea, suggesting that they were likely combined after eukaryogenesis^10^. Our data in this study shows that both RNA-guided RNA cleaving Argonautes and the *rnc-ago-helicase* genomic association exist in one Asgard archaeon. We have not found RdRp in the Asgard archaea. Based on these findings, and combined with the fact that Asgard archaea are the closest known prokaryotic relatives of eukaryotes, we propose a new hypothetical model for the evolutionary origin and diversification of eukaryotic RNA silencing (Fig. 6b). In this model, the eAgo-like RNA-guided RNA cleavage mechanism emerged among the Asgard archaea Argonautes, and formed genomic associations with genes encoding Rnc and DExD/H Helicase, leading to a primordial RNA silencing pathway. Gene rearrangements, duplication, and fusion occurred during the dynamic genome evolution around the period of eukaryogenesis, giving rise to Dicer-like proteins with different types of domain combinations similar to those found in extant eukaryotes. During eukaryotic lineage expansion, dsRNA processing by Dicer was specialized to provide guides for AGOs, while distinct guide-generating pathways, such as the piRNA pathway, developed to provide guides for PIWIs (at least in Metazoa). Structural divergence occurred in adaptations to the specific functions executed by these specialized pathways, such as the loss of g5-g6 seed pre-organization in PIWIs.

The physiological functions of HrAgo1 or other asAgos in their native organisms are yet undetermined due to the inability to cultivate Ago-encoding Asgard archaea. Future *in vivo* studies of eAgo-like asAgos in the context of co-encoded Dicer-domain-containing proteins could shed more light on the mechanisms and functions of these putative RNA silencing pathways. It is possible that asAgos may have fulfilled eAgo-like roles in Asgard archaea, including gene silencing^7,8^, TE silencing^9,58^, maintenance of potential heterochromatins^59^, and/or antiviral defense^60^. Such functionality may have been important in the Asgard archaeal ancestor of eukaryotes to overcome the small genome sizes commonly associated with prokaryotic physiology, and/or in their arms race against mobile genetic elements, enabling a eukaryote-scale genome expansion^13,15,61^.

## Methods

### Identification and selection of Argonautes encoded by Asgard archaea

A custom-built Hidden Markov Model (HMM) encompassing MID-PIWI domain representatives from all known prokaryotic and eukaryotic Argonaute types is provided as a supplementary file (Table S7). This HMM was used to search across 496 Asgard archaea MAGs from NCBI, yielding 138 putative asAgo sequences. Since some sequences are truncated due to fragmented genome assembly, we identified their gene position and the presence of start codon and stop codon to determine the completeness of the genes. The information is provided in Table S1. Incomplete sequences were excluded from phylogenetic analyses except for ASG308_00888, which is the only close homolog of HrAgo1 found in this study but truncated at its N terminus due to contig break.

### Phylogenetic analysis of Asgard archaeal Argonaute

To examine the phylogenetic relation between asAgos and known Argonaute proteins, previously identified pAgos^31^ were first clustered at 60% identity using CD-HIT^62^ v4.8.1. This set was aligned using MAFFT^63^ v7.475 option auto, and sequences with clear N-terminal or C-terminal truncations were removed. The alignment was trimmed using trimAl^64^ v1.4.1 option gappyout, and phylogenetically analyzed using IQtree^65^ v2.1.12 model LG+R9 with 2000 ultrafast bootstrap replicates. The tree was reduced using Treemmer^66^ v0.3 to represent the diversity with fewer related sequences, and well-studied pAgo representatives (highlighted in Fig. 1a) were manually added back if were removed by Treemmer. Next, the well-studied, structurally characterized canonical PIWI and AGO clade proteins were selected to comprise 9 eAgo representatives. The reference Argonaute proteins highlighted in Fig. 1a are PIWI from *Ephydatia flauviatilis* Piwi (EfPIWI), AGO from *Homo sapiens* (hAGO2), archaeal Argonautes from *Pyrococcus furiosus* (PfAgo), *Methanocaldococcus jannaschii* (MjAgo), and *Natronobacterium gregoryi* (NgAgo), *Archaeoglobus fulgidus* (AfAgo), *Sulfolobus islandicus* (SiAgo), and bacterial Argonautes from *Aquifex aeolicus* (AaAgo), *Thermus thermophilus* (TtAgo), *Clostridium butyricum* (CbAgo), *Marinotoga piezophila* (MpAgo), *Rhodobacter sphaeroides* (RsAgo), *Pseudooceanicola lipolyticus* (PliAgo), *Runella slithyformis* (RslAgo), *Crenotalea thermophila* (CrtAgo), *Kordia jejudonensis* (KjAgo), *Xanthomonas vesicatoria* (XavAgo), and *Joostella marina* (JomAgo). 109 asAgos, quality-filtered as described above, were used. The final set comprises a total of 334 Argonautes. These proteins were aligned using MAFFT option linsi, and the MID-PIWI section was retained using the amino acid positions in the HrAgo1 structure as reference. The cropped alignment was then trimmed using trimAl option gt 0.1 to remove the most highly variable regions and used for phylogenetic analysis. Maximum likelihood phylogenetic analysis was carried out using IQtree v2.1.12. The best fitting model was identified using ModelFinder^67^ among all combinations of the LG, WAG, and Q.pfam models combined with the empirical profile mixture model C60^68^, and with modeled rate heterogeneity (either +R4 and +G4). The Q.pfam+C60+F+R4 was selected by the ModelFinder. Statistical support was evaluated using 1,000 replicates via ultrafast boostrap 2 (UFBoot2)^69^. The phylogenetic tree was visualized using iTOL^70^, where ultrafast bootstrap values above 95 were indicated in Fig. 1a.

To examine the stability of the Long-A pAgo branches sister to the eAgo clade, we used two different alignment combinations and three different models. Besides the MID-PIWI domains of all Ago types described above, we omitted the short pAgo clade and made a full-length alignment encompassing the N-L1-PAZ-L2-MID-PIWI domains. In addition to the Q.pfam+C60+F+R4, we also used LG+C60+F+R4 and WAG+C60+F+R4. Statistical support was evaluated using 1,000 replicates via UFBoot2. Branches closest to the eAgo clade were shown in Fig. S2.

Diverse eukaryotic AGO and PIWI full length sequences were used to created HMM profiles via HMMER (http://hmmer.org/). To ensure the full recruitment of evolutionary intermediates between AGO and PIWI, the medium bitscore of AGO members was used as cutoff for PIWI HMM searches, and vice versa. These profiles and bitscore cutoffs were used to recruit eukaryotic Argonaute proteins from the EukProt v3 database^71^. After quality filtering by removing truncated sequences lacking the major domains of Argonaute, 1312 putative AGOs and 454 putative PIWIs were aligned using MAFFT option auto and phylogenetically analysed using FastTree^72^ v2.1.10 model LG. The AGO clade and PIWI clade of the trees were pruned using Treemmer down to 100 branches each, where each eukaryotic supergroup was forced to keep at least 3 sequences if possible. 201 eukaryotic Argonaute representatives were combined with HrAgo1 and ASG308_00888 (the truncated homolog of HrAgo1), TrypAgos, and LongA pAgo sequences, aligned using MAFFT option linsi, trimmed using trimAl option gt 0.1, and analysed using Iqtree v2.1.12. The best fitting model was identified using ModelFinder among all combinations of the LG, WAG, and Q.pfam models combined with the empirical profile mixture model C60, and with modeled rate heterogeneity (either +R4 and +G4). Statistical support was evaluated using 1,000 replicates via UFBoot2. The phylogenetic tree was visualized using iTOL.

### Identification of various features in the ‘*Ca.* H. repetitus’ genome

The present ‘*Ca.* H. repetitus FW102’ genome assembly is a single scaffold with two gaps (GenBank accession: JAIZWK010000001.1). The basic features including the origin of replication protein Cdc6 and 16S and 23S rRNA subunits was annotated as described previously^13^. The CRISPR-Cas operon was annotated using CCTyper^73^. Other defense systems were identified using the Defense-Finder online tool^32^, which also identified HrAgo2 and the CRISPR-Cas system, but did not identify HrAgo1.

### Presence of Argonaute homologs across prokaryotic lineages

The custom MID-PIWI HMM profile was used to search for Argonaute homologs in the GTDB database v207 (for all prokaryotic phyla except Asgard archaea) and an Asgard archaea database (387 genomes after quality filtering using the same standard as GTDB). Prokaryotic phyla with less than 40 representatives were removed for comparison.

### Sequence similarity between HrAgo1 with various eAgos and Long pAgos

Representative sequences were each aligned with HrAgo1, the number of aligned sites with the same identity was divided by the total number of amino acids in HrAgo1 as metrics for sequence similarity.

### Identification of RNaseIII and their genomic association with Argonaute

A custom RNaseIII HMM profile was used for the identification of Rnase III from the Asgard archaea. Potential genomic neighbors, with no more than one gene in-between based on sequence headers were then manually examined at the genomic level. Neighboring genes containing DExD/H helicase domains were identified using Conserved Domain Database^74^.

### Plasmid construction

The *HrAgo1* gene, codon-optimized for *E. coli* and synthesized by Genscript, Inc., was inserted under the T7 promoter in the expression plasmid pET28a to yield pFWC01 (*Pt7::HrAgo1*). A plasmid suitable for expression of a HrAgo1 catalytic double-mutant (D585A & E623A; HrAgo1^DM^) was generated by Quikchange Site-Directed Mutagenesis using primers oPB199 and oPB201 for D585A and oPB200 and oPB198 for E623A, using *E. coli* strain NEB 5-alpha (New England Biolabs) (Table S5).

pX-sfGFP vector was a kind gift from Prof. Jae-Sung Woo (Korea University, South Korea). Linear pX vector backbone was prepared by PCR with primers bypassing sfGFP coding region and then subjected to gel purification. Insert DNA fragments with human codon-optimized coding sequences for FLAG-hAGO2 and FLAG-HrAGO1, flanked by pX vector homology regions, were synthesized commercially (Twist Bioscience). Insert DNA fragments were cloned into the linear pX vector backbone by Gibson assembly (in lab). Competent *E. coli* cells were transformed with the Gibson assembly products, and plasmids (pX-FLAG-hAGO2 and pX-FLAG-HrAGO1) were purified using PureYield Plasmid Miniprep System (Promega).

pmirGLO Dual-Luciferase miRNA Target Expression Vector (Promega) was linearized by PCR with primers that insert two fully complementary binding sites (“perfect target sites”) for human miR-1-1 3p in the 3′ UTR of the firefly luciferase gene. Competent *E. coli* cells were transformed with the linearized vectors, and plasmids (pmirGLO-2X miR-1-1 perfect target site) were purified by miniprep.

pLKO.1 puro was a gift from Bob Weinberg (Addgene plasmid # 8453; http://n2t.net/addgene:8453; RRID:Addgene_8453)^75^. pLKO.1 puro vector was linearized by PCR with primers that insert human pri-mir-1-1 sequence in the downstream of the U6 promoter. Competent *E. coli* cells were transformed with linearized vectors, and plasmids (pLKO.1 puro-pri-mir-1-1) were purified by miniprep. All plasmids were verified by Sanger sequencing (Macrogen). The cloning primers are listed in Table S6.

### HrAgo1 expression and purification

HrAgo1 was heterologously expressed in *Escherichia coli* BL21-Gold (DE3). Expression cultures were shaken at 120 rpm in an incubator at 37°C in LB supplemented with 50 mg/ml kanamycin until an optical density at 600 nm (OD_600 nm_) of 0.4 was reached. The incubation temperature was then decreased to 18°C. When the OD_600 nm_ reached 0.6, expression of HrAgo1 was induced by adding isopropyl-b-D-thiogalactoside (IPTG) to a final concentration of 0.2 mM. Expression of HrAgo1 took place at 18°C for 20 hours. Cells were harvested by centrifugation at 4,000 x g at 4°C for 30 minutes and were lysed by sonication (QSONICA Q700A-220 sonicator with ½” tip, amp 35%, 1s ON/2 s OFF for 4 minutes) in Lysis Buffer (1 M NaCl, 5 mM Imidazole, 20 mM Tris-HCl pH 8) supplemented with protease inhibitors (100 μg/ml AEBSF and 1 μg/ml Pepstatin A). After centrifugation at 40,000 x g at 4°C for 45 minutes, the cell free extract was loaded on 5 ml HisTrap HP column (Cytiva Life Sciences) which was subsequently washed with 25 ml of Washing Buffer I (1 M NaCl, 20 mM Imidazole, 20 mM Tri-HCl pH 8). Bound protein was eluted with Elution Buffer I (1 M NaCl, 250 mM Imidazole, 20 mM Tri-HCl pH 8). The eluted protein was loaded on a custom 20 ml amylose resin column and was washed with Washing Buffer II (1 M NaCl, 20 mM Tri-HCl pH 8, 1 mM DTT). The protein was eluted with Elution Buffer II (1 M NaCl, 20 mM Tri-HCl pH8, 10 mM Maltose, 1 mM DTT). TEV protease was added in a 1:50 (w/w) ratio (TEV:total protein), and the mixture was dialyzed overnight in SnakeSkin dialysis tubing (30kDa MWCO, Thermo Scientific) against 2l dialysis buffer (1M KCl, 20 mM HEPES-KOH pH 7.5, 1 mM DTT, 2 mM EDTA) at 4°C for 16 h. TEV-mediated removal of the His-MBP tag was confirmed by SDS-PAGE analysis. The sample was concentrated to a volume of 1 ml using 30 K centrifugal filter units (Amicon). After concentrating, the sample was centrifuged for 10 min at 16,000 x g at 4°C to remove aggregates and the supernatant was loaded on a custom 200 ml Superdex 200 resin column which was pre-equilibrated with SEC buffer (1 M KCl, 20 mM HEPES-KOH pH 7.5, 1 mM DTT). The peak fractions were analysed by SDS-PAGE and fractions containing HrAgo1 were combined and concentrated, aliquoted and flash frozen in liquid nitrogen before storage at −70°C until further use. HrAgo1^DM^ was expressed and purified as HrAgo1 with minor modifications: For expression *E. coli* BL21 Star (DE3) was used. Furthermore, expression was performed in TB medium containing 20 µg/ml kanamycin.

### Cleavage activity assays

HrAgo1 activity assays were performed in reactions with a final volume of 20 µl with the following final concentrations: 0.4 µM HrAgo1, 0.4 µM guide oligonucleotide (ogDS001, ogDS002, ogDS003, or oBK458 (Table S2)), 0.1 µM Cy5-labelled target oligonucleotide (oDS401 or oDS403; Table S2), 5 mM HEPES-KOH, 125 mM KCl, and 2 mM divalent metal salt (MnCl_2_ or MgCl_2_)). Prior to addition of the target, HrAgo1 and the guide were incubated for 15 min at 37°C. After addition of the target, HrAgo1:guide:target ratios were 4:4:1. The mixture was incubated for 1h at 37°C. The reaction was stopped by adding 2X RNA Loading Dye (250 mM EDTA, 5% v/v glycerol, 95% v/v formamide) and further incubation at 95°C for 10 min. The samples were resolved on a 20% denaturing (7 M Urea) polyacrylamide gel. The gels were imaged on an Ettan DIGE Imager (GE Healthcare (480/530 nm)). Time-dependent cleavage assays were performed in a similar way but with a HrAgo1:guide:target ratio of 4:2:1.

### Small RNA extraction and analysis

Two nanomoles of purified HrAgo1 were incubated with 250 µg/ml Proteinase K (Thermo Scientific) for 4 h at 65°C. Next, phenol:chloroform:IAA 25:24:1 pH 7.9 (Invitrogen) was added in a 1:1 ratio. The sample was vortexed and centrifuged at 16000 x g in a table top centrifuge for 10 min. The upper layer containing the nucleic acids was transferred to a clean tube and the nucleic acids were precipitated through ethanol precipitation. To this end, 99% cold ethanol and 3 M sodium acetate pH 5.2 were added to the sample in a 2:1 and 1:9 ratio, respectively. The sample was incubated overnight at −80°C, after which it was centrifuged at 16000 x g in a table top centrifuge for 1 h. The pellet was washed with 70% ethanol and subsequently dissolved in nuclease-free water.

Purified nucleic acids were [γ-^32^P]-ATP labelled with T4 polynucleotide kinase (PNK; Thermo Scientific) in an exchange-labelling reaction. After stopping the reaction by incubation at 75 °C for 10 min, the labelled oligonucleotides were separated from free [γ-^32^P] ATP using a custom Sephadex G-25 column (GE Healthcare). Labelled nucleic acids were incubated with nucleases (Rnase A, Dnase and protease-free (Thermo Scientific), or Dnase I, Rnase-free (Thermo Scientific) for 30 min at 37 °C. After nuclease treatment, samples were mixed with Loading Buffer (95% (deionized) formamide, 5 mM EDTA, 0.025% SDS, 0.025% bromophenol blue and 0.025% xylene cyanol), heated for 5 min at 95 °C and resolved on 15% denaturing (7M Urea) polyacrylamide gels. Radioactivity was captured from gels using phosphor screens and imaged using a Typhoon FLA 7000 laser-scanner, GE Healthcare).

Small RNA sequencing libraries were prepared and sequenced by GenomeScan (Leiden, The Netherlands) using Illumina NovaSeq6000 sequencing with paired-end reads and 150bp read length. Paired-end small RNA reads were merged, adapter sequences were trimmed, and length was trimmed to 35 nucleotides using Bbtools v38.90^76^. Processed reads of all sequencing libraries were aligned to the genome of *E. coli* BL21 (GenBank: CP053602.1) and to the expression plasmid (pFWC01) using HISAT2 v2.1.0^77^. Length, sequence distribution, and abundance of specific small RNAs were analysed using FastQC (https://www.bioinformatics.babraham.ac.uk/projects/fastqc/) after extracting uniquely mapped reads using HISAT2 and Samtools v1.2^78^.

### Cryo-EM sample preparation and data collection

Purified HrAgo1 was mixed with a 5’-phosphorylated RNA guide (5’-UGAGGUAGUAGGUUGUAUAGU-3’) in assembly buffer (5 mM HEPES pH 7.5, 250 mM KCl, 5 mM MgCl_2_). The final sample contained 8.6 μM HrAgo1 and 8.6 μM of g-RNA in a total volume of 60 μL. The volume was incubated at 37 °C for 15 minutes and centrifuged at 18000 rpm for 10 min at room temperature. After adding CHAPSO (Sigma-Aldrich) to a final concentration of 0.8 mM, the sample was used for cryo-EM grid preparation. 2.5 µL of the above sample was applied to a freshly glow discharged 300-mesh UltrAuF R1.2/1.3 grid (Quantifoil Micro Tools), blotted for 5 s at 100% humidity, 4 °C, plunge frozen in liquid ethane (using a Vitrobot Mark IV plunger, FEI) and stored in liquid nitrogen. Cryo-EM data collection was performed on a FEI Titan Krios G3i microscope (University of Zurich, Switzerland) operated at 300 kV and equipped with a Gatan K3 direct electron detector in super-resolution counting mode. A total of 8977 movies were recorded at 130000x magnification, resulting in a super-resolution pixel size of 0.325 Å. Each movie comprised 47 subframes with a total dose of 56.81 e^-^/Å^2^. Data acquisition was performed with EPU Automated Data Acquisition Software for Single Particle Analysis (ThermoFisher Scientific) with three shots per hole at −1.0 mm to −2.4 mm defocus (0.2 mm steps).

### CryoEM data processing and model building

The collected exposures were processed in cryoSPARC (v.4.2)^79^. Patch Motion Correction and Patch CTF Correction were used to align and correct the imported 8977 movies. Movies with CTF resolution higher than 20 Å were discarded, resulting in a total of accepted 8275 movies. Template picker (particle diameter 140 Å; templates were selected from a previous data collection on the same sample) was used to select particles, which were included for further processing based on their NCC and power score. Particles were extracted (extraction box size 360 pix; Fourier-cropped to box size 120 pix) and classified in 50 classes using 2D Classification. 22 classes (2188198 particles) were selected and given as input to a 2-classes Ab-Initio Reconstruction. The 1299949 particles corresponding to one of the two reconstructions were further sorted in 100 classes using 2D Classification. 28 classes (533275 particles) were used for a 2-classes Ab-Initio Reconstruction (maximum resolution 6 Å; initial resolution 20 Å; initial minibatch size 300; final minibatch size 2000). The particles of one of the two reconstructions were assigned to 80 classes using 2D classification, 57 of which (283659 particles) were extracted to full resolution and selected for non-uniform refinement (initial lowpass resolution 20 Å; per-particle CTF parameters and defocus optimization). A final round of non-uniform refinement (dynamic mask start resolution 1 Å; initial lowpass resolution 20 Å; per-particle CTF parameters and defocus optimization) resulted in a 3.40 Å (GSFSC resolution, FSC cutoff 0.143) density. A detailed processing workflow is shown in the Supplementary Figure S4.

An initial model of HrAgo1 was generated using AlphaFold2 ColabFold^80^. The model was manually docked as rigid body in the cryoEM density map using UCSF ChimeraX^81^, followed by real space fitting with the Fit in Map function. The model was subjected to manual refinement against the corresponding cryoEM map using the software Coot^82^ and real space refine in Phenix^83^. Secondary structure restraints, side chain rotamer restraints and Ramachandran restraints were used. The final model comprises one copy of HrAgo1(27-99,103-193,198-271,282-307,322-589,595-817), one copy of the gRNA (1-6) and 1 Mg^2+^ ion. Low resolution density for the RNA 3’ end was visible in the map, but not confidently interpretable, therefore it was not built in the final model. Figures preparation of model and map was performed using UCSF ChimeraX.

### Single-molecule experimental set-up

All single-molecule experiments were performed on a custom-built microscope setup. An inverted microscope (IX73, Olympus) with prism-based total internal reflection was used in combination with a 532 nm diode-pumped solid-state laser (Compass 215M/50mW, Coherent). Photons are collected with a 60x water immersion objective (UPLSAPO60XW, Olympus), after which a 532 nm long pass filter (LDP01-532RU-25, Semrock) blocks the excitation light. A dichroic mirror (635 dcxr, Chroma) separates the fluorescence signal which is then projected onto an EM-CCD camera (iXon Ultra, DU-897U-CS0-#BV, Andor Technology).

### Single-molecule sample preparation

Synthetic RNA was purchased from Horizon Discovery (United Kingdom). The guide and target strands (Table S4) were labelled with Cy5 Mono NHS Ester and Cy3 Mono NHS Ester (Sigma-Aldrich), respectively. 5 µl of 200 µM RNA, 1 µl of 0.5 M freshly prepared 0.5 M sodium bicarbonate and 1 µl of 20 mM dye in DMSO were mixed and incubated overnight at 4°C in the dark, followed by ethanol precipitation. The labeling efficiency was ∼100%. The target strands were subsequently ligated with a biotinylated polyuridine strand (U_30_-biotin). To this end, 200 pmol of target RNA strand was mixed with U_30_-biotin and a DNA splint in a 1:1:3 ratio in TE buffer with 100 mM NaCl. The mixture was annealed in a thermal cycler by rapidly heating it to 80 °C for 4 min and then slowly cooling it down with 1 °C every 4 min. The annealed constructs were ligated using 2 µL T4 RNA ligase2 (NEB, 10U/µL), 3 µL 0.1% BSA (Ambion), 3 µL 10x reaction buffer (NEB), 0.25 µl 1 M MgCl_2_ and 0.3 µl Rnasin ribonuclease inhibitor (Promega, 0.4 u/µl) in a final volume of 30 µl at 25°C overnight. After acidic phenol-chloroform extraction and ethanol precipitation, the ligated RNA strands were purified on a 10% denaturing (7M urea) polyacrylamide gel.

For the t1-target assays, the RNA target strands were produced through in vitro transcription of DNA templates (Table S4). All synthetic DNA was purchased from Ella Biotech (Germany). First, an annealing mix was prepared with template DNA and IVT T7 promotor oligos at a final concentration of 40 µM each in a 10 µl reaction with 1x annealing buffer (50 mM NaCl and 10 mM Tris-HCl pH 8.0). The annealing mix was heated to 90 °C for 3 min and then slowly cooled with 1 °C every min to 4 °C. Next, in vitro transcription was performed using the TranscriptAid T7 High Yield kit (Thermo Scientific) for 4 hours at 37 °C according to the manufacturer’s instructions. After acidic phenol-chloroform extraction and ethanol precipitation, the RNA strands were purified on a 10% denaturing (7M urea) polyacrylamide gel. Finally, the purified RNA target strands (2 µM in 10 µl) were annealed to the immobilization strand and imager strand in a 2:1:5 ratio in annealing buffer by heating to 90 °C for 3 min and then slowly cooling with 1 °C every min to 4 °C.

Microfluidic chambers with a polymer(PEG)-coated quartz surface were prepared as described previously^84^. Each chamber was incubated with 20 µl 0.1 mg/ml Streptavidin (Sigma) for 30 s. Unbound Streptavidin was flushed out with 100 µl T50 (10 mM Tris-HCl pH 8.0, 50 mM NaCl). Next, 50 µl 50 pM Cy3-labeled target RNA was introduced into the chamber and incubated for 1 min. Unbound target RNA was flushed out with 100 µl T50 and 100 µl imaging buffer (50 mM Tris-HCl pH 8.0, 500 mM NaCl, 1 mM Trolox (Sigma), 0.8% glucose, 0.5 mg/mL glucose oxidase (Sigma), 85 µg/mL catalase (Merck), 0.4 u/µl Rnasin ribonuclease inhibitor (Promega)) was introduced into the chamber. EfPiwi was purified as previously described^5^. For EfPiwi, binding was much weaker so to enable observation of these events within our time resolution 50 mM instead of 500 mM NaCl was used in the imaging buffer. The PIWI binary complex was formed by incubating 15 nM purified PIWI in imaging buffer (minus the glucose oxidase and catalase which were added after incubation) with 1 nM Cy5-labeled guide RNA at 37°C for 10 min. The binary complex was introduced in the chamber, after which 200 s long movies were recorded. The experiments were performed at room temperature (22 ± 2 °C).

### Single-molecule data acquisition and analysis

CCD movies of time resolution 0.1 s were acquired using Andor Solis software v4.32. Co-localization between the Cy3 and Cy5 signal and time trace extraction were carried out using Python. The extracted time traces were processed using FRETboard v0.0.3^85^. The dissociation rate was estimated by measuring the dwell times off all binding events. The dwell time distributions were fit with an exponential decay curve (Ae^-t/Δτ^) or with the sum of two exponential decay curves (A1e^-t/Δτ1^+ A2e^-t/Δτ2^).

### Mammalian cell culture and transfection

HCT116 *AGO1/2/3* knockout (KO) cells were obtained from the Corey lab (UT Southwestern, USA). Cells were grown and maintained in McCoy’s 5A Modified Medium (Thermo Fisher Scientific) supplemented with 9% (v/v) fetal bovine serum (Cytiva) in an incubator at 37°C and 5% CO_2_.

For stable transfection, 2E6 KO cells were seeded in a 9 ml medium on a 100 mm culture dish 1 day before transfection. Transfection was performed with 5 µg of pLKO.1 puro pri-mir-1-1 vector using Lipofectamine 3000 (Thermo Fisher Scientific) according to the manufacturer’s instructions. 2 days after the transfection, culture medium was replaced by a medium containing 2 µg/ml puromycin (Thermo Fisher Scientific) (“selection medium”). The selection proceeded for 16 days, and the selection medium was replaced every 4 days.

For transient transfection, 2E6 puromycin-selected cells were seeded in a 9 ml medium on a 100 mm culture dish 1 day before transfection. The cells were co-transfected with 4 µg of pX vector (pX-sfGFP or pX-FLAG-hAGO2 or pX-FLAG-HrAGO1) and 1 µg of pmirGLO-2X miR-1-1 perfect target site using Lipofectamine 3000. Cells were harvested 2 days after transfection, snap frozen by liquid nitrogen, and then stored at −80°C.

### RT-qPCR

Total RNAs were isolated using TRIzol (Thermo Fisher Scientific), treated with RQ1 RNase-Free DNase (Promega), and then phenol-extracted. Complementary DNAs (cDNAs) were synthesized from 2 µg of total RNAs using SuperScript IV Reverse Transcriptase (Thermo Fisher Scientific) and random hexamer according to the manufacturer’s instructions. qPCR was performed using Power SYBR Green PCR Master Mix (Thermo Fisher Scientific) and QuantStudio 5 Real-Time PCR systems. The qPCR primers are listed in Table S6.

### Western blotting

Cells were lysed by re-suspension in ice-cold lysis buffer (150 mM NaCl, 50 mM Tris-HCl pH 7.5, 1% Triton X-100, 0.5% Sodium Deoxycholate, 0.1% SDS) supplemented with Protease Inhibitor Cocktail Set III, EDTA-Free (Millipore). The lysed cells were centrifuged at 16,100 x g, 4°C, 15 min, and the supernatant (“total protein lysate”) was transferred to a fresh e-tube. The concentration of the total protein lysates was measured by Pierce BCA Assay (Thermo Fisher Scientific). 50 µg of total protein lysates were boiled with 4x Laemmli Sample Buffer (Bio-Rad), run on NuPAGE 4-12% Bis-Tris gel (Thermo Fisher Scientific) with PageRuler Plus Prestained Protein Ladder (Thermo Fisher Scientific), and then transferred onto methanol-activated Immun-Blot PVDF membrane (Bio-Rad) using Mini Blot Module (Thermo Fisher Scientific). The membrane was blocked in PBS-T (PBS (PanReac AppliChem) + 0.1% Tween-20 (Sigma)) containing 5% skim milk, probed with primary antibodies at 4°C, ON, and then washed three times with PBS-T. Rabbit polyclonal FLAG antibody (1:1000, Sigma, F7425) was used to probe ectopically expressed FLAG-hAGO2 or FLAG-HrAGO1, and rat monoclonal Tubulin antibody (1:1000, Invitrogen, MA1-80017) was used to probe loading control. The washed membranes were probed with secondary antibodies at RT for 1 h, and then washed three times with PBS-T. Alexa Fluor 647-conjugated donkey anti-rabbit IgG (1:2000, Jackson ImmunoResearch, 711-605-152) and Alexa Fluor 546-conjugated goat anti-rat IgG (1:2000, Invitrogen, A-11081) were used as secondary antibodies. The protein bands were detected by fluorescence using the Typhoon laser-scanner platform system (Cytiva).

## Data availability

HMM profiles, protein sequence alignment files, phylogenetic trees, and CryoEM structure and validation report listed in Table S7 and will be provided upon publication of the manuscript. The small RNA sequencing data will be made available at the Gene Expression Omnibus database upon publication of the manuscript. Atomic coordinates and cryo-EM maps have been deposited in the protein data bank (PDB entry ID 8R3Z) and Electron Microscopy Data Bank (EMDB, entry ID EMD-18878) and will be made public upon publication of the manuscript.

## Acknowledgements

HCT116 knockout cells were a kind gift from Prof. David R. Corey (UT Southwestern, USA). F.W. thanks Woodward Fischer and Diaoqiong Zheng for valuable discussions, and Danxi Cui for technical support. C.B. and C.J. thank Martin Depken and Hidde Offerhaus for valuable discussions on the data analysis. F.W. was supported by National Science Foundation of China grant (32370003). C.J. was supported by ERC Consolidator grant (819299) of the European Research Council. D.C.S was supported by grants from the European Research Council (ERC-2020-STG 948783) and Veni grant (016.Veni.192.072). P.B.U. was supported by Consejo Nacional de Ciencia y Technology (CVU No. 682509). K.K. was supported by an EMBO Postdoctoral Fellowship (ALTF 76-2022).

## Contributions

F.W. conceived the project and supervised it with C.J. and D.C.S.. F.W. and Y.F. performed protein identification and phylogenetic analyses. P.B.U. and F.W. constructed plasmids and purified HrAgo1 proteins. T.A.A. purified EfPiwi proteins. C.B. isolated and sequenced small RNAs. D.C.S. performed HrAgo1-associated small RNA analyses. P.B.U. performed *in vitro* cleavage assays. C.B. performed single-molecule assay and analyzed the data with C.J.. G.F. performed CryoEM and analyzed data with D.C.S. and I.J.M.. K.K. constructed plasmids and performed RNA silencing experiments in human cell lines. S.K., D.T., and T.J.G.E. provided additional asAgo sequences and phylogenetic pipelines. F.W., D.C.S., C.J., C.B., P.B.U., K.K., G.F., and I.J.M. wrote the manuscript. All authors commented on the manuscript.

## Supplementary Figures

**Fig. S1.**
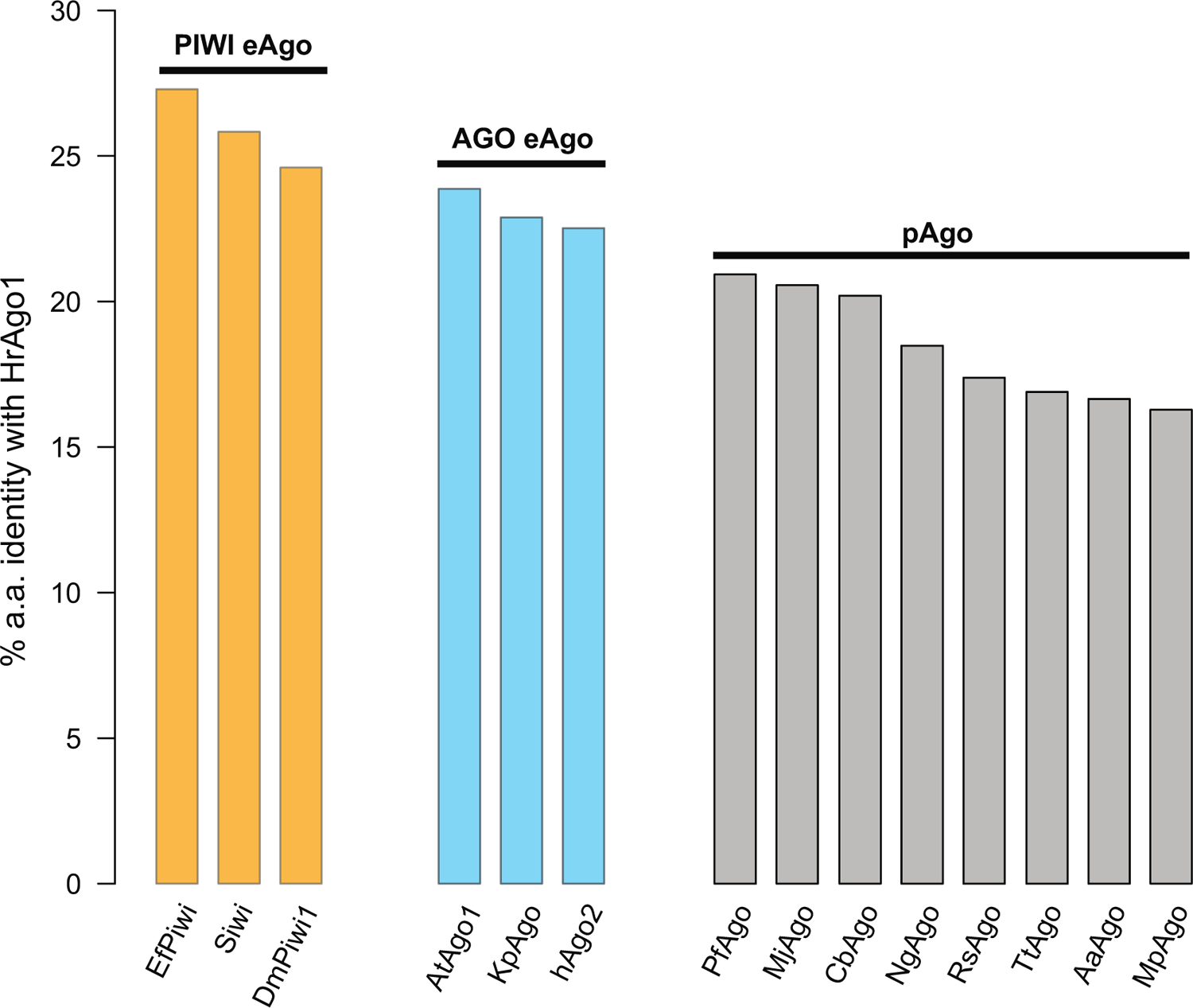
Percentage amino acid identity conservation between HrAgo1 and various biochemically studied pAgos and eAgos.

**Fig. S2.**
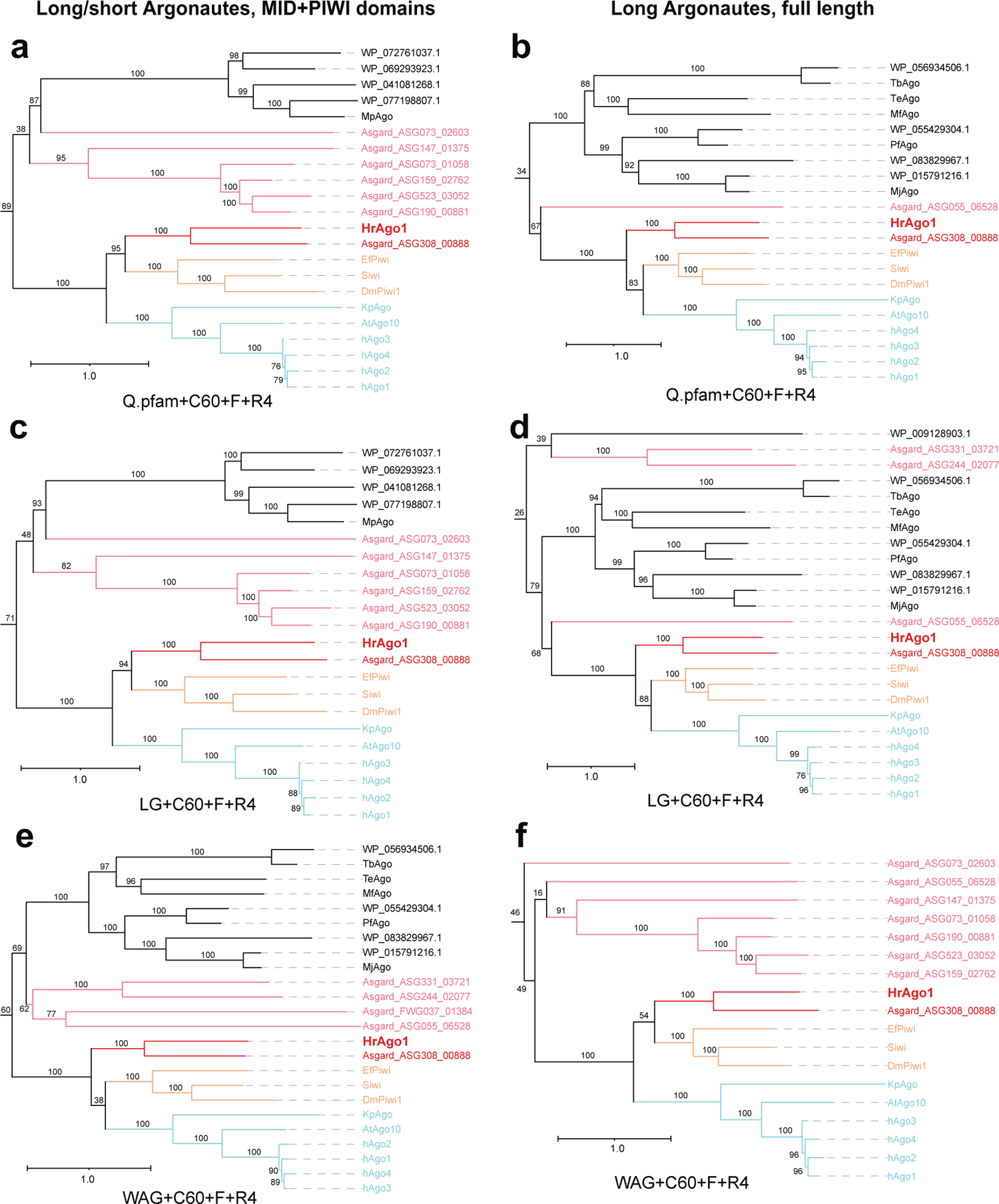
Maximum-likelihood pAgo-asAgo-eAgo phylogenetic analyses support the close relation between HrAgo1 and eAgos, while the exact root of the eAgo clade is unstable. The analyses were done with the MID+PIWI domains of all Ago types (Left), and the full-length alignment of Long-A, Long-B, and the HrAgo2 clade (right), using three different kinds of mixture models under 1000 ultrafast bootstrap replicates in IQtree. Only branches close to the eAgo clade is shown. In a, c, and f, HrAgo1 is sister to the PIWI clade, while in b, d, and e, HrAgo1 is sister to the whole eAgo clade. This reflects a basal position that is difficult to resolve. Some other asAgos (in pink) appeared basal to the eAgo-HrAgo1 clade in multiple conditions, but are supported by low bootstrap values. UFBoot2 values are shown.

**Fig. S3.**
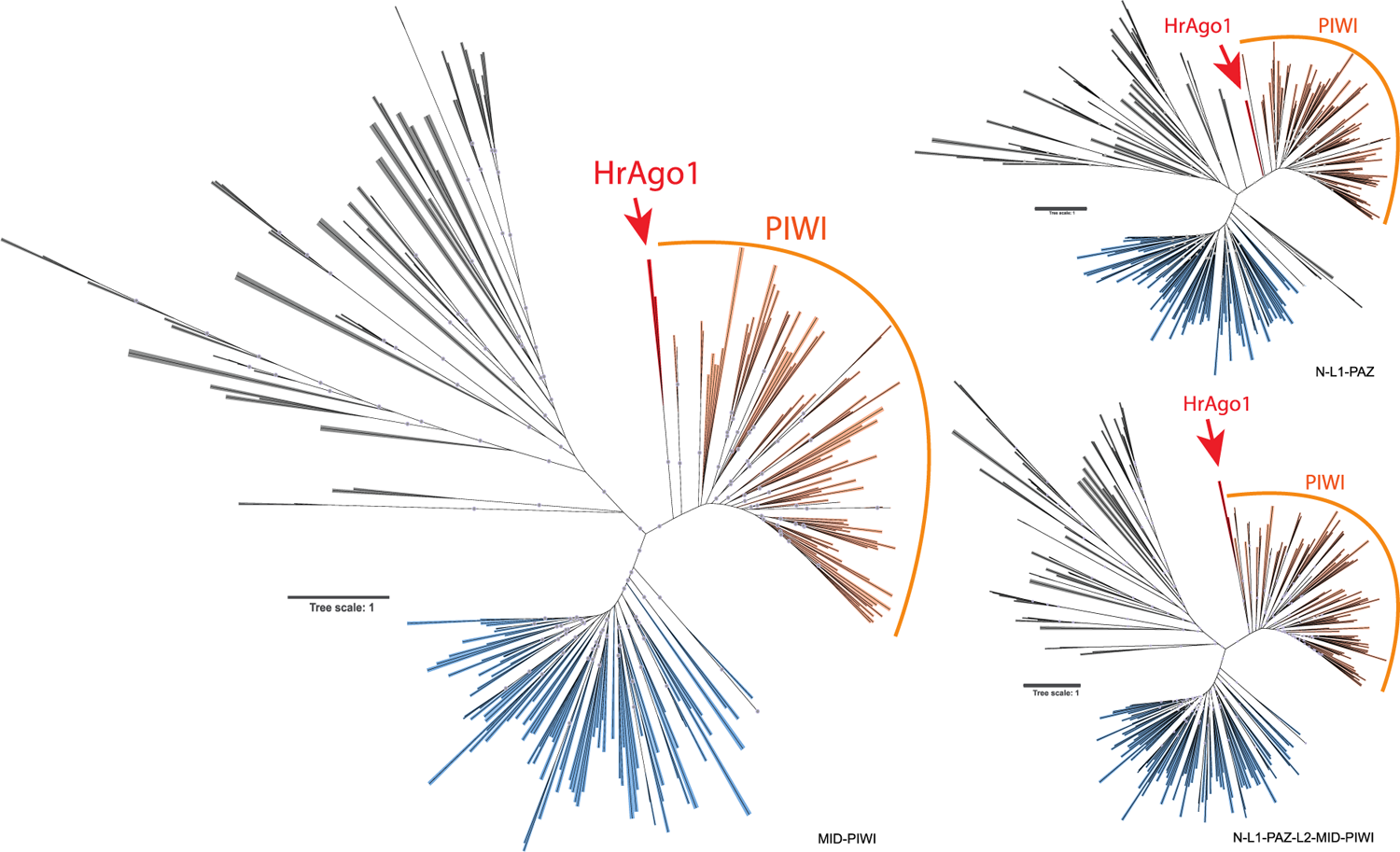
Unrooted phylogenetic trees of different protein domain combinations showing that HrAgo1 is consistently positioned basal to the PIWI clade. Grey, pAgo and TrypAgos. Blue, PIWI clade. Orange, AGO clade. Red, HrAgo1 clade. The domains used for phylogenetic analyses are indicated at the bottom right of each tree. Grey circles indicate UFBoot2 values above 90.

**Fig. S4.**
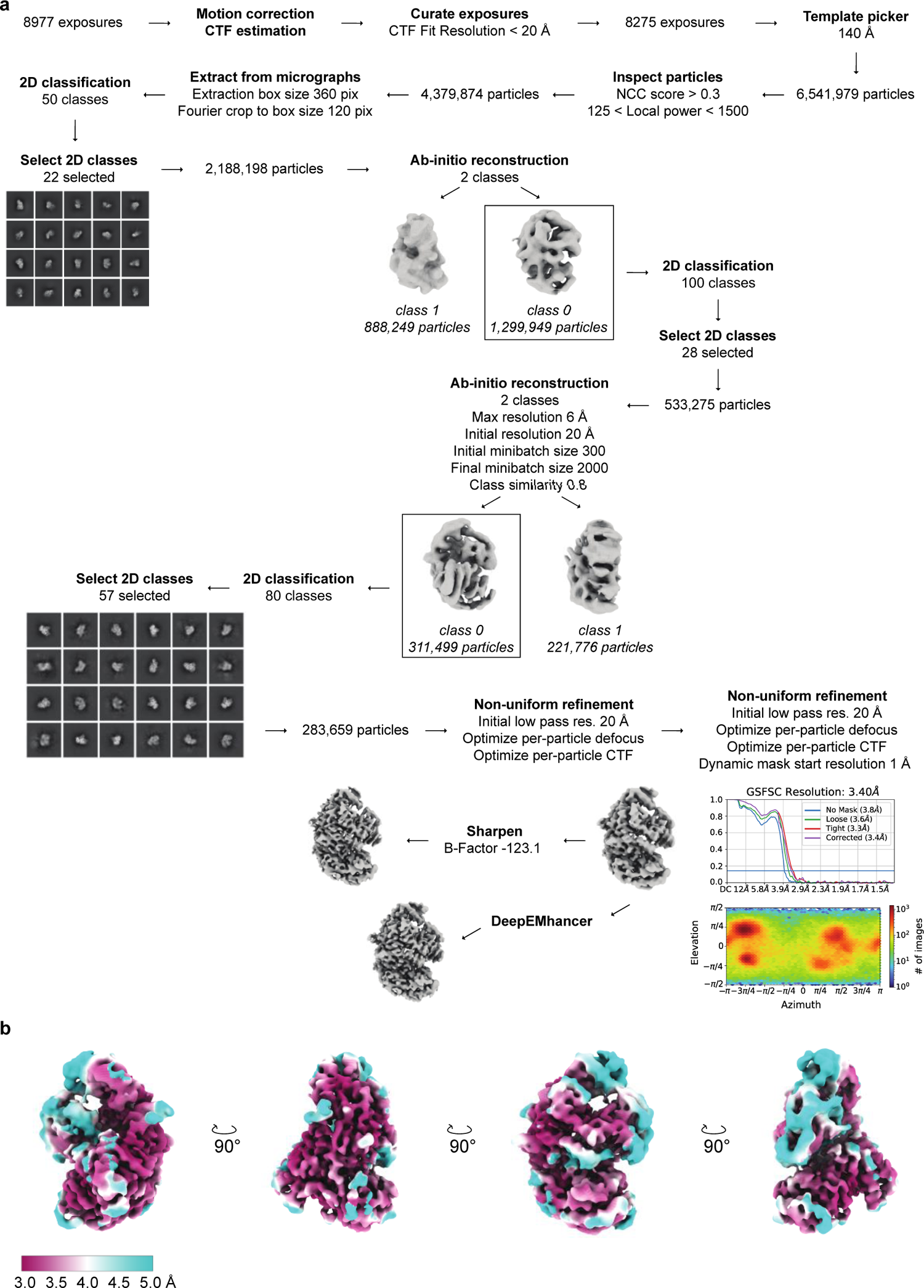
Cryo-EM data processing for HrAgo1-gRNA complex. **a**, Cryo-EM image processing workflow for HrAgo1-gRNA. Unless specified, standard processing parameters were used. **b,** Cryo-EM densities of the HrAgo1-gRNA complex colored according to local resolution.

**Fig. S5.**
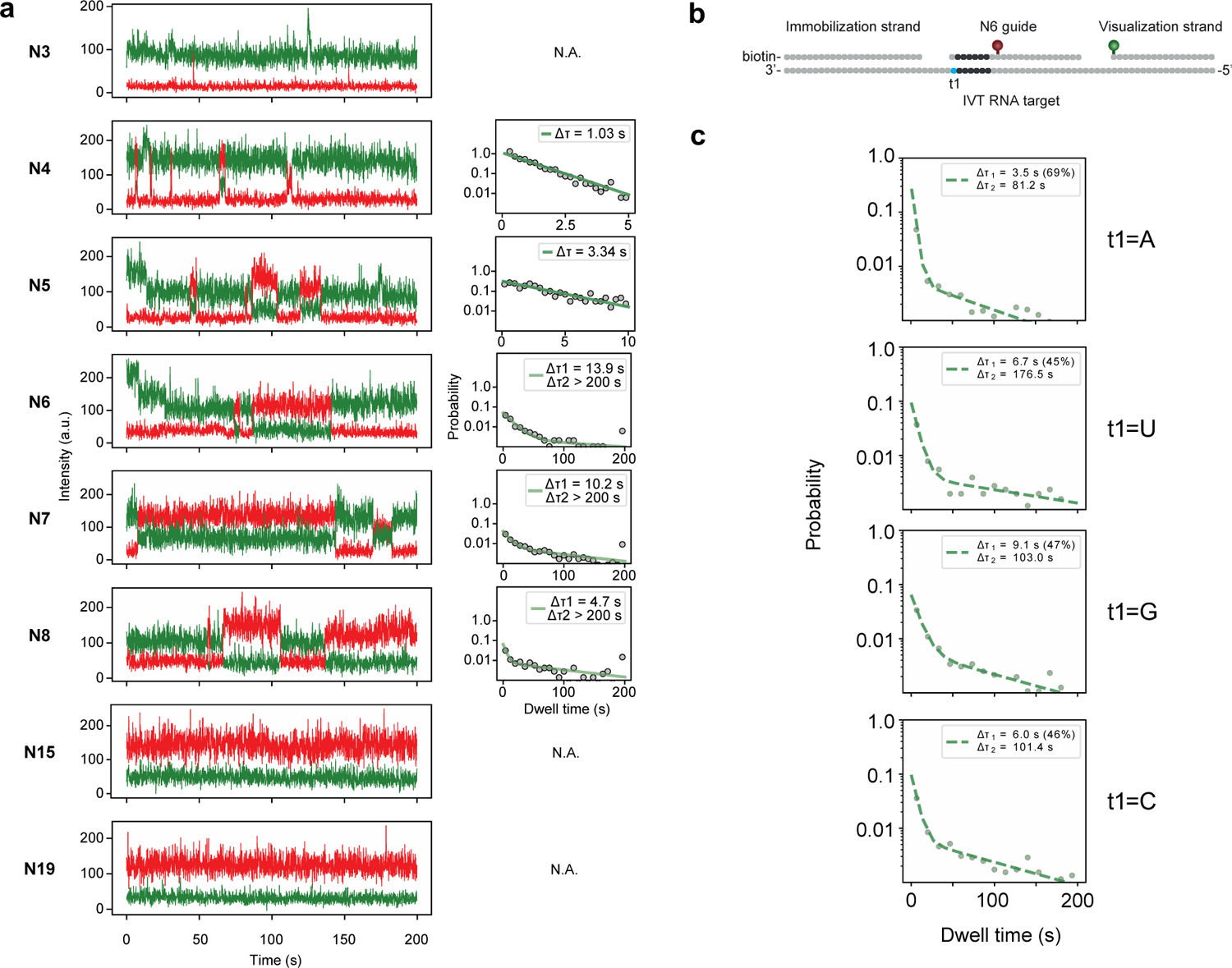
Representative time traces and dwell time distributions with fit for HrAgo1. **a**, Representative time traces (left) and dwell time distributions with fit (right) for HrAgo1. Dwell time distributions were fit with a single or double exponential. N, match length between guide and target starting from the second nucleotide. N.A., no fit due to time resolution or observation time limit. Due to the observation time limit, the second dwell time is underestimated, therefore it is set to > 200 s for all match lengths that exhibit a stably bound population. For N3, the time resolution is limiting so the dwell time is set to < 0.1 s. **b**, Schematic of the construct used for the t1-target assays. **c,** Dwell time distributions with double exponential fit for targets with a different nucleotide at the first position (t1).

**Fig. S6.**
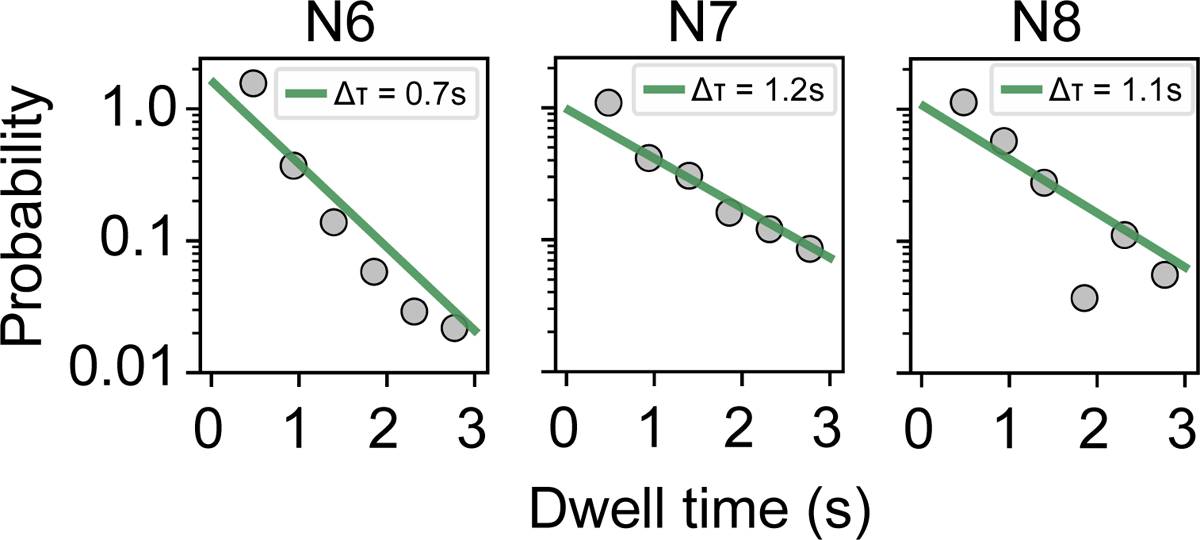
Dwell time distributions with fit for EfPiwi. The dwell time distributions were fit with a single exponential. For shorter and longer match lengths, the time resolution (0.1s) and observation time (200s) were limiting so no fit could be made.

**Fig. S7.**
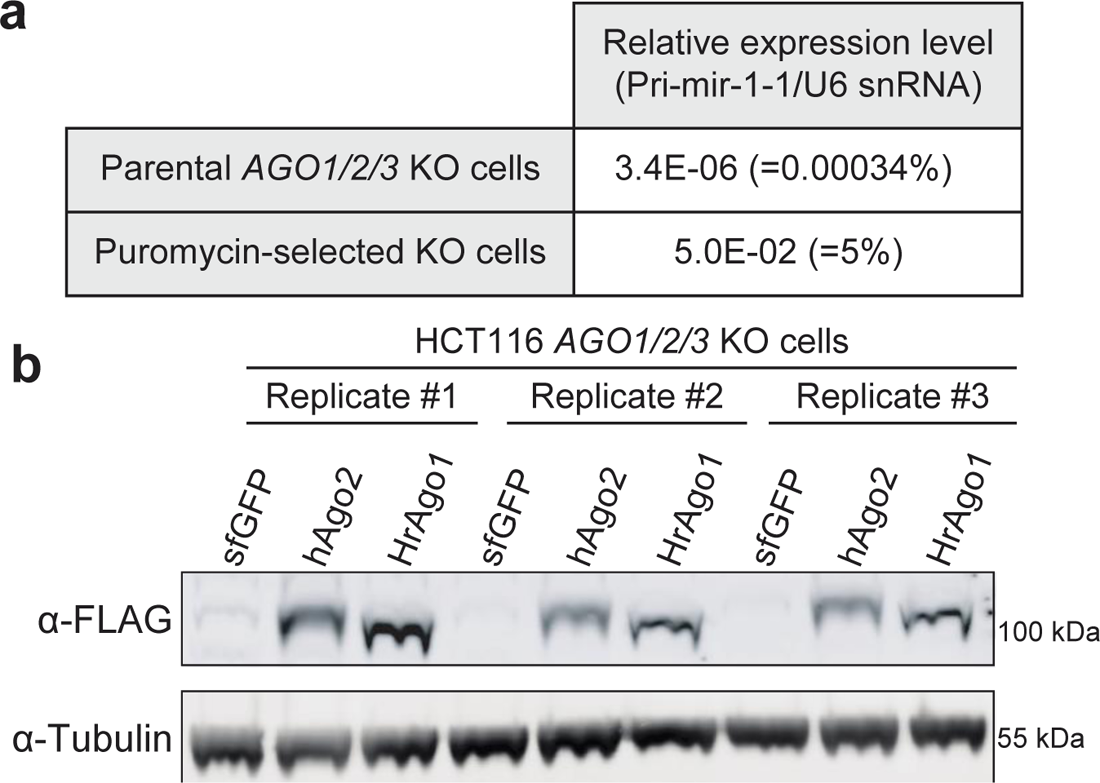
Characterization of engineered human cell lines. **a**, qPCR results for pri-mir-1-1 from parental cells and puromycin-selected cells. Expression levels were normalized to U6 snRNA. **b,** Western blot results for ectopically expressed FLAG-hAGO2 and FLAG-HrAGO1. Tubulin was used as loading control.

